# Structural Basis of Prostaglandin Efflux by MRP4

**DOI:** 10.1101/2022.12.22.521501

**Authors:** Sergei Pourmal, Evan Green, Ruchika Bajaj, Ilan E. Chemmama, Giselle M. Knudsen, Meghna Gupta, Andrej Sali, Yifan Cheng, Charles S. Craik, Deanna L. Kroetz, Robert M. Stroud

**Affiliations:** Department of Biochemistry and Biophysics, University of California, San Francisco, CA; Department of Chemistry and Chemical Biology Graduate Program, University of California, San Francisco, CA; Department of Biophysics Graduate Program, University of California, San Francisco, CA; Department of Bioengineering and Therapeutic Sciences, University of California, San Francisco, CA; Department of Pharmaceutical Chemistry, University of California, San Francisco, CA; Quantitative Biosciences Institute, University of California, San Francisco, CA; Howard Hughes Medical Institute, University of California, San Francisco, CA

## Abstract

MRP4 is unique among the C family of ATP-binding cassette transporters for its role in translocating prostanoids, an important group of signaling molecules derived from unsaturated fatty acids. Using a reconstituted system, we report that a pair of prostaglandins (PGs) and the sulfonated-sterol DHEA-S preferentially enhance the ATPase activity of MRP4 over other previously proposed physiological substrates such as cyclic nucleotides or leukotrienes. We determined the cryo-EM structures of nanodisc embedded bovine MRP4 in (i) a nucleotide- and substrate-free state, (ii) in complex with PGE_1_, (iii) PGE_2_, and (iv) DHEA-S, and (v) a catalytically dead mutant E1202Q bound to ATP-Mg^2+^. The substrate-bound structures suggest unique features of the MRP4 binding site that distinguish its specificity for prostanoids from that of the related leukotriene transporter MRP1. The ATP-bound structure is in an outward-occluded conformation, revealing a novel state in the proposed alternate-access mechanism of MRP transport. Our study provides insights into the endogenous function of this versatile efflux transporter.

## Introduction

The prostanoid family of lipid mediators are important signaling molecules involved in physiological processes as diverse as inflammation, nociception, immune response, vasoactivity, and parturition^1–7^. Prostanoids are derived from the cyclooxygenase (COX1/COX2)-mediated metabolism of unsaturated fatty acids and subsequently converted in a tissue-specific manner into various prostaglandins (PGs) or thromboxanes (TXs)^8^. Acting in an autocrine or paracrine fashion, secreted prostanoids activate their cognate G-protein coupled receptors, leading to modulation of intracellular Ca^2+^ or cAMP levels that affect downstream signaling and gene expression. Altered prostanoid signaling has implications for a variety of pathologies, including thrombosis, blood pressure, gastrointestinal cancers and regulation of the tumor microenvironment^9–11^. Synthetic prostaglandins have clinical use for induction of labor and abortions, treatment of glaucoma and pulmonary hypertension, vasodilation, and the prevention of stomach ulcers^12–16^. Despite the numerous physiological roles of prostanoids and the therapeutic potential of their analogs, structural details on their active transport from the cell are limited.

The efflux of prostaglandins is mediated by the multidrug resistance associated protein 4 (MRP4)^17^. Broadly expressed in tissues including the liver, kidneys, blood-brain barrier, and blood cells, MRP4 is a membrane bound ATP-binding cassette (ABC) transporter from the ABCC family that functions as a unidirectional cellular efflux pump. It is a single polypeptide chain organized as two half transporters in tandem, each consisting of a transmembrane domain (TMD) composed of six transmembrane (TM) helices followed by a nucleotide binding domain (NBD) (Fig. 1a)^18^. It is the only member of the MRP subfamily that has an established role in the transport of prostanoids, including PGE_1_, PGE_2_, PGD_2_, PGF_2α_, and TXB_2_^17,19,20^. Among these substrates, PGE_2_ is of interest as the most abundant prostanoid produced by the cyclooxygenases and for its suggested roles in cancer pathology. PGE_2_ promotes tumor cell growth and metastasis through the activation of E-prostanoid (EP) receptors and the transactivation of growth factor signaling. It further facilitates cancer aggressiveness by suppressing anti-tumor immunity while enhancing tumoral immune evasion^9,11,21^. The requirement for PGs to be exported for EP signaling, the rapid rate of PG metabolism in the cytoplasm, and the lack of any alternative known PG exporter implicate MRP4 in pathological PG efflux. Consistent with this role for MRP4, its overexpression in breast tumors has been associated with poor prognosis^22^, suggesting therapeutic potential in modulating MRP4 function.

**Fig 1.**
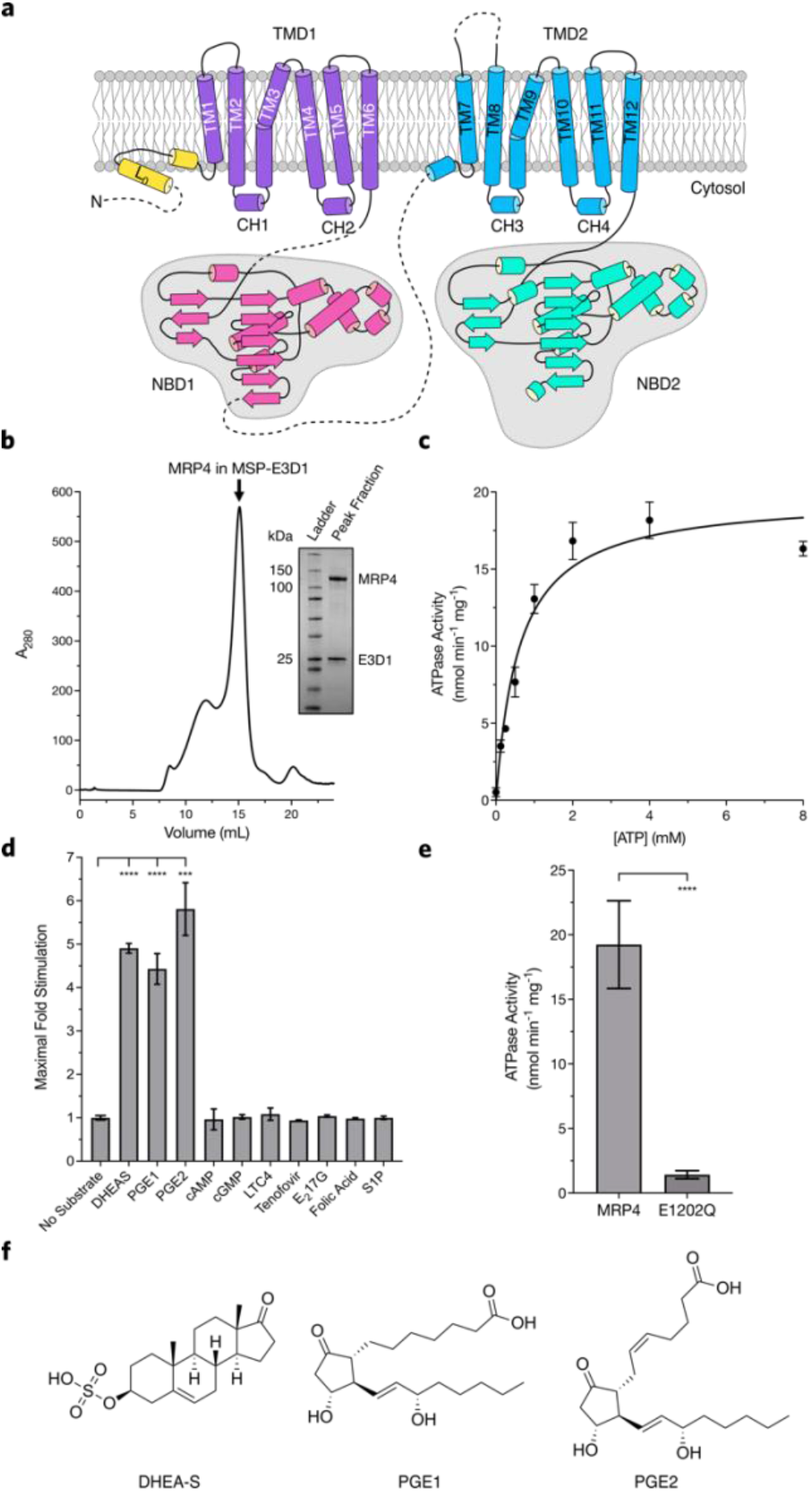
Functional characterization of MRP4 and identification of three stimulating substrates. **a**, Schematic illustrating the domain architecture of MRP4 with residue numbers at domain boundaries indicated. L_0_, N-terminal loop domain (yellow); TMD1, transmembrane domain 1 (purple); NBD1, nucleotide-binding domain 1 (pink); TMD2, transmembrane domain 2 (blue); NBD2, nucleotide-binding domain 2 (cyan). **b**, Gel-filtration chromatography profile of WT MRP4 embedded into MSP-E3D1 lipid nanodiscs. SDS-PAGE analysis of peak fraction is inserted on the right, visualized by Coomassie blue staining. **c**, ATPase activity of nanodisc embedded MRP4. Data shown are the mean ± SD for 3 technical replicates and fit to the Michaelis-Menten equation. Source data are provided as a Source Data file. **d**, WT MRP4 ATPase activity with 4 mM ATP in the presence of previously reported substrates. ATPase activity shown is for the highest concentration tested for a given substrate, normalized to MRP4’s basal activity without a substrate. Titration curves at different concentrations of a given substrate and the effects on ATPase activity are in Supplementary Data Fig 1a-f. Data shown are mean ± SD for 3 technical replicates. Source data are provided as a Source Data file. **e**, Comparison of WT MRP4 and MRP4_EQ_ ATPase activity in lipid nanodiscs. Data shown are mean ± SD for 3 technical replicates. Source data are provided as a Source Data file. **f**, Structures of previously reported transport substrates found to stimulate MRP4 ATPase activity.

In addition to its role in PG efflux, MRP4 has been linked to the transport of numerous structurally diverse substrates. MRP4 overlaps with other MRP family members in the transport of steroid conjugates, including dehydroepiandrosterone sulfate (DHEA-S), the steroid with the highest concentrations in human circulation and a precursor for androgens and estrogens, and estradiol 17β-D-glucuronide (E217βG), a conjugated metabolite of estradiol^23^. MRP4 has also been classified as a cyclic nucleotide transporter^24^. Studies in *Mrp4-/-* mice have linked cyclic nucleotide transport by MRP4 to vascular reactivity and resistance to hypoxic pulmonary hypertension^25^, impaired platelet activation^26^, and cystic fibrosis transmembrane conductance regulator-mediated chloride flux and secretory diarrhea^27,28^. Other substrates of MRP4 identified through overexpression systems include cysteinyl leukotrienes, bile salts, folate, sphingosine-1-phosphate (S1P), cephalosporin antibiotics, and antivirals^23,29–37^, though to date none of these substrates, including the prostanoids, DHEA-S and E217βG, have been validated in reconstitution experiments.

Structural information on MRP4 and its substrate binding has long been limited to homology-based models. More recently, AlphaFold2 and other protein folding software have provided atomic-level predictions, though these models are currently limited to a single state of this dynamic transporter. In order to better understand the molecular details of MRP4 function and to identify the structural elements that define MRP4 substrate selectivity, we used cryo-electron microscopy (cryo-EM) to determine high-resolution structures of lipid nanodisc-reconstituted MRP4 under five conditions. Our nucleotide- and substrate-free (apo-), three substrate-bound (DHEA-S, PGE_1_ and PGE_2_) and ATP-bound structures represent three distinct states along the MRP4 substrate translocation cycle. Along with our *in vitro* examination of MRP4 ATPase activity, the substrate-bound structures provide insights into the basis of substrate discrimination among closely related members of the MRP subfamily. Our ATP-bound MRP4 structure expands our understanding of the ABCC family conformational landscape by revealing the outward-facing occluded state, an important intermediate in the proposed alternating access mechanism of substrate translocation. This series of structures will enable further computational investigations into the role of MRP4 in transporting xenobiotic and endogenous substrates and provides a basis for designing small molecules to modulate MRP4-mediated efflux.

## Results

### Purification, nanodisc reconstitution, and functional characterization of MRP4

GFP-tagged orthologs of MRP4 were screened using fluorescence size exclusion chromatography (FSEC). *Bos taurus* MRP4, which shares 90% similarity to human MRP4 (Supplementary Data Fig. 1), was selected for its expression level and biochemical behavior and is the focus of this study. This ortholog was heterologously expressed in *Spodoptera frugiperda* (Sf9) cells with a cleavable C-terminal 8-His tag that allowed for detergent purification using metal-affinity TALON resin followed by SEC (Supplementary Fig. 2a). To pursue ATPase activity assays and cryoEM structure determination in a native lipid environment, we reconstituted MRP4 into MSP E3D1 lipid nanodiscs directly on affinity resin, which substantially increased the efficiency of incorporation into nanodiscs (Fig. 1b).

As Type IV exporters display substrate-dependent increases in ATPase activity ^38–40^, we used stimulation of MRP4 ATP hydrolysis to assess previously identified substrates for cryoEM analysis. The ATPase activity of MRP4 in the absence of any substrate was determined to have a K_m_ for ATP of 0.60 mM (95% confidence interval (CI) 0.47 - 0.77 mM) and V_max_ of 19.7 nmol min^-1^ mg^-1^ (95% CI 18.4 - 21.1 nmol min^-1^ mg^-1^) (Fig. 1c), similar to previously reported values for human MRP4 ^41^; these values correspond to a baseline specific activity of 4.1 min^-1^ (95% CI 3.8 – 4.4 min^-1^) assuming a total molecular weight of 210 kDa for MRP4 and two copies of MSP E3D1. PGE_1_, PGE_2_ and DHEA-S (Fig. 1f) stimulated the ATPase activity of MRP4 in a concentration-dependent manner (Supplementary Data Fig. 1b and 1c; Supplementary Data Table 2), with maximal stimulation of approximately 4-fold for PGE_1_, and 5-fold for PGE_2_ and DHEA-S (Fig. 1d). In contrast, we observed no concentration-dependent stimulation of ATPase activity for several other previously reported MRP4 substrates, including the cyclic nucleotides cAMP and cGMP, the nucleoside analog tenofovir, the sterol E217βG, the vitamin folic acid, the sphingolipid S1P, and the MRP1 substrate leukotriene C_4_ (LTC_4_)^18,42^ (Fig 1d; Supplementary Data Fig. 1d-1g). As the results with PGE_1_, PGE_2_, and DHEA-S suggested the presence of a high-affinity transporter-cargo complex, we pursued structures of MRP4 bound to these three ATPase stimulating substrates.

As typically seen in other Type IV transporters, ATP-binding induces dimerization of the NBDs. The two ATPase sites are composed of residues contributed by both NBDs, as the Walker A and B motifs from one NBD form a site with the signature sequence from the other. In MRP4, sequence analysis (Supplementary Fig. 1) reveals that the NBD2 Walker B motif contributes a catalytic E1202 to form a canonical active ATPase site. In contrast, the NBD1 Walker B contains D560 in the corresponding position, suggesting this latter ATPase site is degenerate and catalytically compromised. In order to pursue a nucleotide-bound structure of MRP4, we introduced an E1202Q mutation (MRP4_E1202Q_) to prevent hydrolysis of ATP while retaining the ability to bind nucleotides. MRP4_E1202Q_ had negligible ATPase activity, consistent with the critical role of E1202 in ATP hydrolysis and confirming the degeneracy of the NBD1 Walker B motif (Fig. 1e).

### Structures of MRP4 in apo, substrate-bound and ATP-bound states determined by cryo-EM

Purified MRP4 in lipid nanodiscs was frozen in the absence of any substrate or nucleotide (apo) directly on cryo-EM grids for data collection. The resulting density map has a resolution of 3.1 Å by the Fourier shell correlation (FSC, 0.143) criterion (Fig. 2a, Supplemental Data Fig. 2, Supplemental Table 1) and has well-defined density for all transmembrane helices and NBD2, which allowed the interpretation of these regions in context of the primary structure. While NBD1 is not defined at near-atomic resolution in the apo density, the domain is well resolved in the subsequent ATP-bound dataset from this study. NBD1 from the ATP-bound state was used as a starting model for positioning and refinement into the apo density. N-terminal segments of the lasso motif and the linker between NBD1 and TMD2 are unresolved.

**Fig 2.**
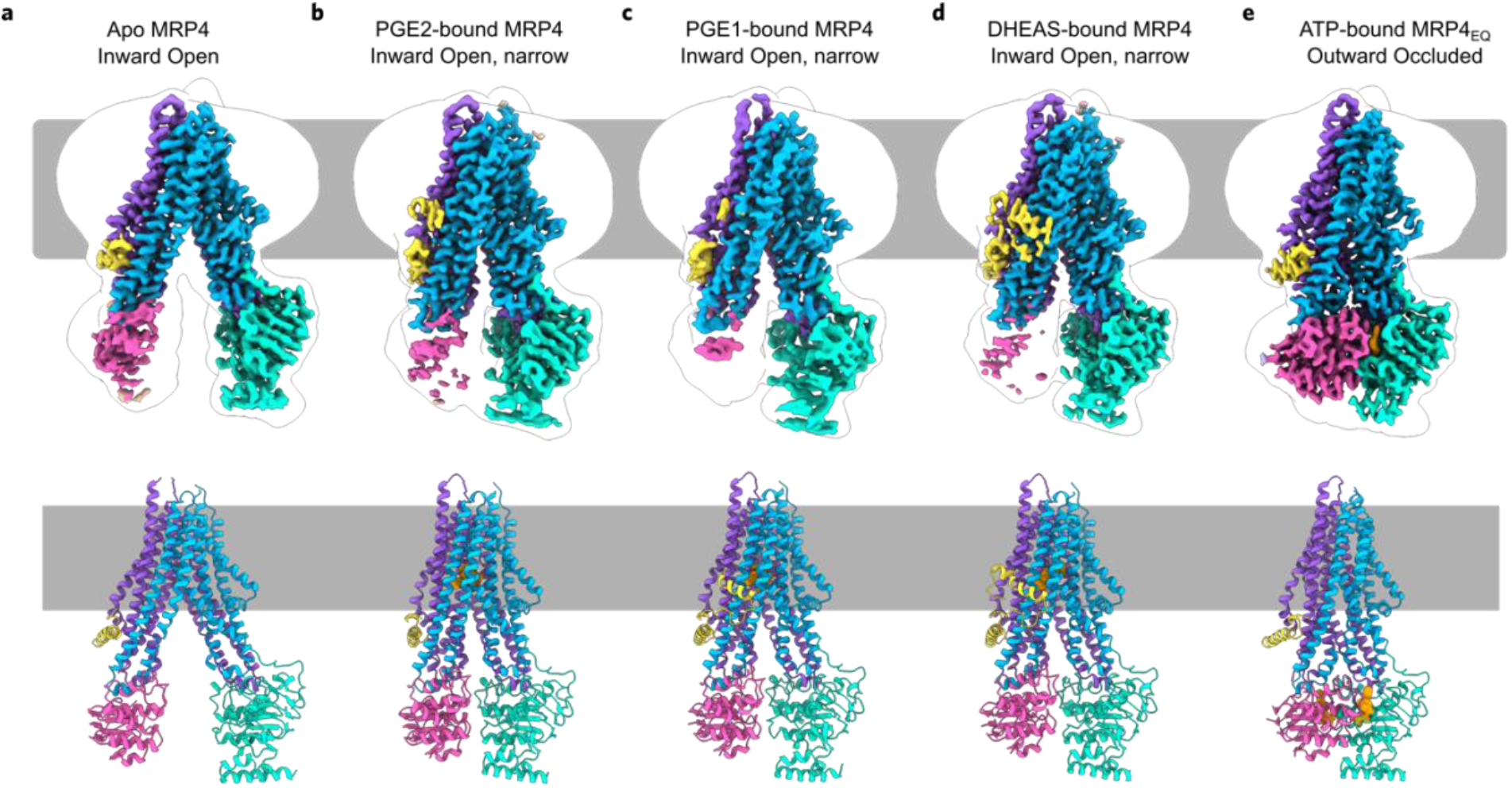
Structures of MRP4 in multiple conformations along the substrate transport cycle. Side views of the cryo-EM reconstruction and atomic model of **a**, inward open, apo-conformation of MRP4, **b**, inward open narrow, DHEA-S bound MRP4 **c**, inward open narrow, PGE1 bound MRP4, **d**, inward open narrow, PGE_2_ bound MRP4, and **e**, outward occluded, ATP-Mg2+ bound MRP4_E1202Q_. The sharpened electrostatic potential maps shown were generated using DeepEMhancer. Domains in sharpened maps and models are colored as in Figure 1a. A transparent low-pass filtered envelope depicts density for the nanodisc and poorly resolved NBDs. Membrane boundaries are represented by cartoon. DHEA-S, PGE1, PGE2, and ATP-Mg^2+^ are shown as orange spheres in their respective structures.

To capture substrate-bound states of MRP4, separate datasets were collected for nanodisc reconstituted MRP4 in the presence of either DHEA-S, PGE_1_ or PGE_2_. The data collected from these complexes yielded reconstructions at 2.7 Å, 3.5 Å and 2.9 Å resolution, respectively (Figs. 2b-2d, Supplementary Data Fig. 3-5, Supplementary Data Table 1), with clear densities for the TMDs, NBD2, and the substrates in their binding sites. Portions of the N-terminal lasso motif, a common feature of ABCC family members that precedes the elbow helix of TMD1, are resolved in the DHEA-S and PGE_1_–bound structures. NBD1 was poorly resolved in all three substrate-bound maps, though the pose and relative orientation could be modeled following the procedure described for the apo structure.

In addition to these three stimulating substrates, a dataset was collected for MRP4 in the presence of cAMP, a substrate which failed to enhance MRP4 ATPase activity *in vitro* yet whose transport has been reported to be MRP4-dependent in various cell types and membrane-derived vesicle assays^24,43^. Cryo-EM analysis of MRP4 in the presence of 1 mM cAMP (concentrations far higher than previously reported Km values and cytoplasmic levels of this important signaling molecule) revealed a structure at 3.7 Å, with no apparent conformational change relative to apo MRP4 (Supplemental Fig. 8a,b) and no strong additional density observed for cAMP in the substrate binding pocket (Supplemental Fig. 8c). The structure of MRP4 in this condition was not modeled.

A final image dataset was collected under saturating ATP-Mg^2+^ conditions, with substrate added to promote progression through the transport cycle. MRP4_E1202Q_ in the presence of nucleotide and PGE_2_ resulted in a 3.1 Å resolution map with high resolution throughout the transporter, including both NBDs bound to ATP and magnesium (Fig. 2e, Supplementary Data Fig. 6, Supplementary Data Table 1). The NBD models built into this map were used to fit NBDs into the apo and substrate-bound states. All five models were refined into their respective densities, and together reveal three distinct conformations of MRP4 at high resolution, including the first structure of an ABCC family exporter in an inward closed, outward-occluded state (Fig. 2e).

The overall architecture of MRP4 resembles that of other mammalian ABC transporters, with two halves each consisting of a TMD and a cytoplasmic NBD, on a single polypeptide chain (Fig. 1a). An N-terminal lasso motif consists of a membrane-embedded helix that packs against TM helices 3, 10 and 11 before continuing into a second extended helix that runs parallel to the membrane’s inner leaflet. The TMD-NBD halves are configured in a pseudo-2-fold symmetric arrangement perpendicular to the membrane, with an unstructured, flexible 82-residue linker connecting NBD1 and TMD2; this linker remains unresolved in all five structures. Similar to other type IV ABC exporters, a pair of TM helices are domain swapped from one TMD to the other, so that helices 1, 2, 3, 6, 10 and 11 form one distinct transmembrane bundle, and TM helices 4, 5, 7, 8, 9 and 12 form another (Fig. 2a-2e). This helical swapping permits extended cytosolic loops in either TMD to interact with both NBD1 and NBD2, allowing for allosteric coupling between soluble and membrane-bound regions. The NBDs of MRP4 each contain a RecA-type core featuring the highly conserved Walker A, Walker B and signature sequence motifs that coordinate the binding and hydrolysis of ATP (Supplementary Fig. 1). Immediately following NBD2 is a 27-residue, unstructured C-terminal linker containing the terminal PDZ-binding sequence ‘ETAL’, which interacts with PDZ adaptor proteins to modulate cell surface expression of MRP4 ^44^.

### Conformational changes upon substrate and ATP binding

The apo state of MRP4 is in an inward open conformation. A solvent-filled cavity (~11,000 Å^3^) is present between the two transmembrane bundles, extending from the cytoplasm into the lipid bilayer, ending at the interface of the bundles and the site of substrate binding. The bundles are at their maximal observed separation and the substrate binding site is accessible, characterized by a 17.1 Å distance between the backbone Cα of a pair of substrate-binding residues, F211 from TMD1 and W995 from TMD2 (Fig. 3a). The NBDs are oriented in a quasi-symmetric head-to-tail fashion and are separated from one another by about 27 Å at either of the two ATPase sites (Fig. 3b), as measured by the Cα distance between the signature sequence serine of one NBD and the Walker A glycine of the other NBD.

**Fig 3.**
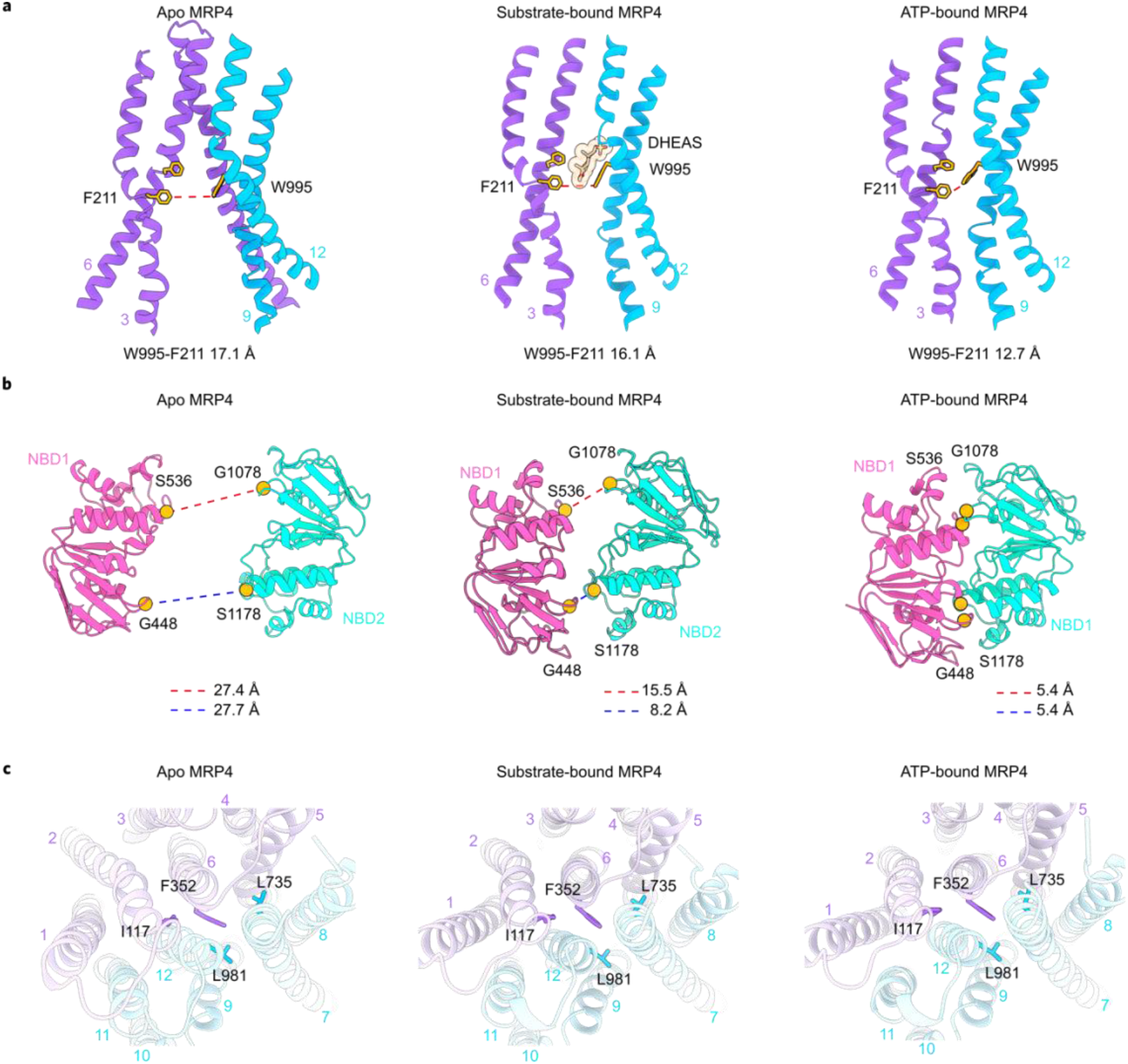
Comparison of the inward open, inward narrow, and outward occluded conformations of MRP4. **a**, Side view of the TMDs from the plane of the membrane for apo-, substrate-bound (DHEA-S bound structure shown), and ATP-bound states. A pair of substrate-binding residues from either TMD is shown in stick representation. The distance between these residues is drawn as a red dashed line, with values indicted in each panel. **b**, Bottom view of the NBDs, seen from the direction of the cytoplasm. Cα of signature sequence serines and Walker A glycines shown as orange spheres. Distances between opposing pairs of residues are shown as dashed lines; red for the functional consensus ATPase site, and blue for the degenerate ATPase site. Distance values are indicted in each panel. **c**, Top down views of the extracellular face. Extracellular gate residues are shown as stick representation. Domains follow the color scheme presented in Figure 1a.

MRP4 undergoes a clamshell-like closure upon substrate binding. The angle between the two transmembrane bundles decreases by 20°, leaving the central cavity more constricted (~5,400 Å^3^) compared to the apo state. Though this conformation is narrower, substrate-bound MRP4 remains open toward the cytoplasm. At the substrate-binding site, the distance between F211 and W995 is reduced by 1 Å, indicating only modest rearrangements in the transmembrane region (Fig. 3a). More pronounced conformational changes are observed in the distal cytoplasmic portions of MRP4, where the two NBDs move closer together and are now slightly askew. The narrowing of the transporter results in unequal distances of 8 and 15 Å between the Walker A of one NBD and signature sequence of the other NBD (Fig 3b). Binding of any of the three ATPase stimulating substrates results in similar changes in MRP4. The largest RMSD among the three substrate-bound structures is 1.2 Å, suggesting they represent a common substrate-bound conformation (Supplementary Fig. 9).

The structure of MRP4_E1202Q_ in the presence of PGE_2_ and ATP exhibits substantial global changes compared to apo and substrate-bound MRP4. The NBDs dimerize around two copies of ATP-Mg^2+^ and the transmembrane bundles tightly pack in a parallel orientation, resulting in a conformation that is closed to the interior of the cell (Fig. 3a and 3b). The cytoplasmic portions of TM3 and TM10 contact TM4 and TM9 respectively, forming an intracellular gate that removes access to the central cavity. The movement of the transmembrane bundles occludes the substrate binding site, as the backbones of F211 and W995 move 4 Å closer and their sidechains make van der Waals contacts, preventing substrate binding. This closure of MRP4’s soluble cavity follows transport of substrate across the membrane, consistent with the lack of observed PGE_2_ density in this state.

The extracellular surface of MRP4 remains closed and is highly superimposable across all states (Fig. 3c). Hydrophobic residues I117, F352, L735 and L981 pack together to form an extracellular gate that seals the interface of the two TMDs. The presence of a closed gate in our ATP-bound structure suggests that the substrate exit pathway at the extracellular face can reset independent of ATP hydrolysis and NBD dissociation, possibly to maintain unidirectional transport. Throughout MRP4’s substrate transport cycle, the two halves of the transporter move as rigid bodies around a pivot point at the extracellular surface and show only minor conformational rearrangements within the pair of TMDs and NBDs (Supplementary Data Fig. 8).

### Structural features of NBDs and nucleotide binding

The interface between the TMDs and NBDs is essential for connecting substrate transport to ATP hydrolysis. As in other type IV ABC exporters, a pair of coupling helices (CH) in the loops between the domain-swapped TMs 4,5 and 10,11 fit into a shallow cleft in either NBD (Fig. 4b and 4c). Compared to NBD1, NBD2 contains an insertion of 13 residues between the Walker A motif and the signature sequence (Fig. 4a). These residues (1099-1111) form two short helices that cap the distal end of the NBD2 cleft and tightly pack with the linker between TM12 and NBD2 (Fig. 4c). H1111 from the second inserted helix hydrogen bonds with E292 from TM5 while simultaneously forming π-π interactions with W1025 from the TM12-NBD2 linker. A turn of the helix away, R1114 forms a salt bridge with E1022, further stabilizing contact with the linker (Fig 4d). The absence of a corresponding region in NBD1 results in a smaller interface and fewer contacts with the cytoplasmic extensions of transmembrane helices (Fig. 4b), possibly contributing to the lower resolution of NBD1 density observed across apo- and all substrate-bound states (Supplementary Data Figs. 3f, 4f, 5f, 6f & 7f).

**Fig 4.**
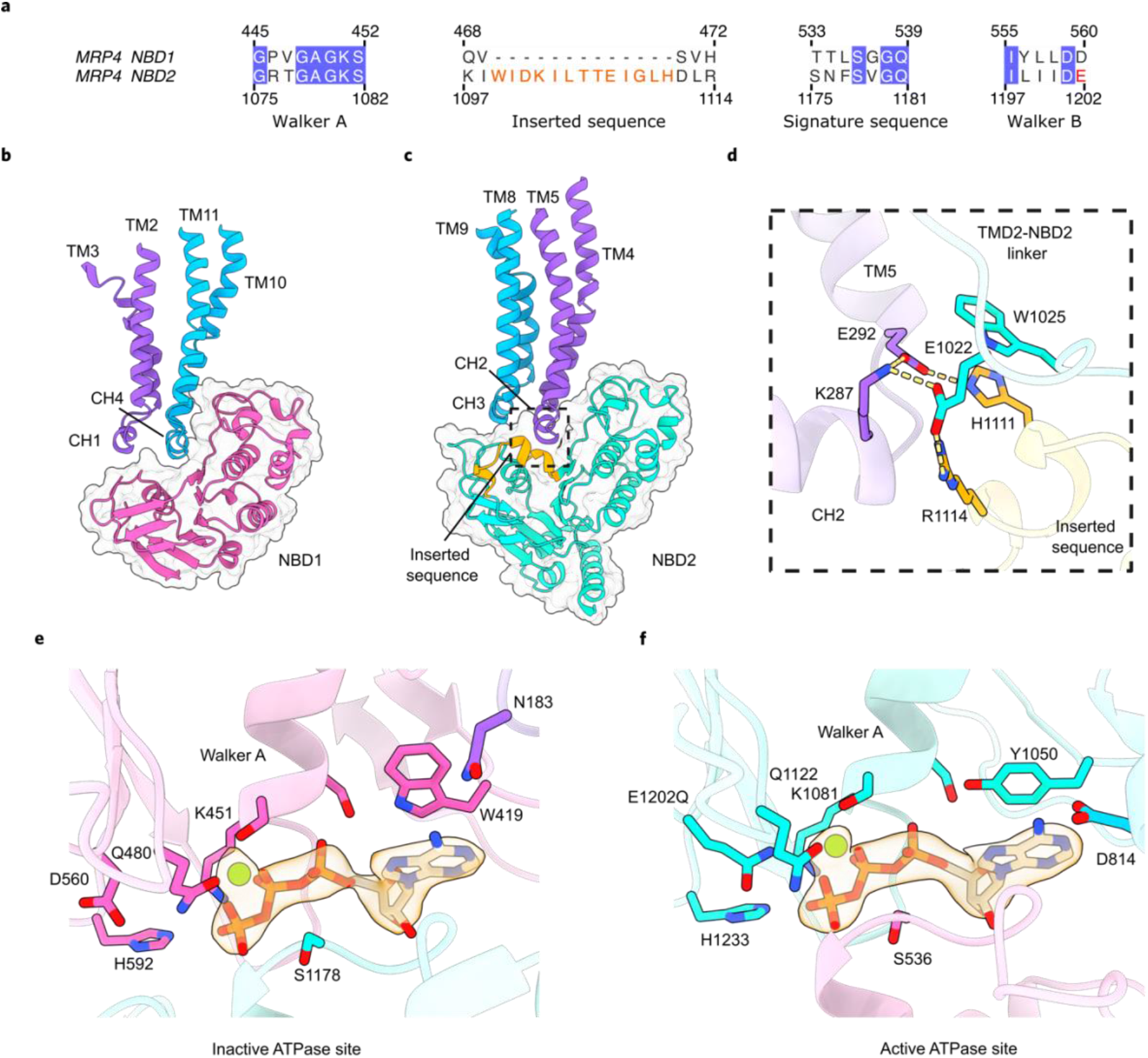
Structural asymmetry between NBD1 and NBD2. **a**, Sequence alignment of selected motifs from NBD1 and NBD2. Conserved residues are highlighted in blue, residues of the inserted sequence are orange, and the position of E1202Q mutant in NBD2 Walker B is in red. Side view of the TMD interactions with **b**, NBD1 and **c**, NBD2. Ribbon diagram of domains are colored as in Figure 1a, and are overlaid on a surface of either NBD in transparent grey. The inserted sequence in NBD2 is highlighted in orange. **d**, Enlarged view of the box in **c**. Electrostatic interactions between the inserted sequence, CH2, and the TMD2-NBD2 linker, with residues shown as sticks. Hydrogen bonds and salt bridges are indicated by yellow dashed lines. Close-up views of the **e**, degenerate and **f**, functional ATPase sites in the ATP-Mg^2+^-bound, outward occluded state. Selected side chains from interacting motifs are shown as sticks. ATP is shown as sticks, Mg^2+^ as green spheres, and densities for both are shown as a transparent surface. For clarity, only ATP-Mg^2+^ density is shown. Selected electrostatic interactions are shown as yellow dashed lines.

In the ATP-bound state, clear density defines the ATP and Mg^2+^ in the E1202Q mutated consensus site, and in the degenerate ATP binding site (Supplementary Fig. 7b). ATP binding is mediated in both sites by highly conserved features. These include electrostatic interactions between Mg^2+^ and the Walker A motif (K451, K1081), and between phosphates and the Walker B motif (D560, E1202Q), the Q loop (Q480, Q1122) and the His loop (H592, H1233) from one NBD and the signature sequence (S536, S1178) from the other NBD (Fig. 4e and 4f). Aromatic residues (W419, Y1050) coordinate the adenine rings of ATP through π-π interactions (Fig. 4e and 4f). The distance of the catalytic carboxyl group of D560 from the phosphorus of the γ-phosphate of ATP of 4.6 Å in the E1202Q mutant is increased to 6.1 Å in the inactive degenerate site. This increased separation accounts for the inability of D560 to catalyze ATP hydrolysis at the degenerate site. Residues from CH1 and CH3 are also important in coordinating ATP binding. Specifically, N183 and D814 coordinate the NH2 group of the adenine ring in the degenerate and consensus site, respectively (Fig. 4e and 4f).

### Substrate binding by MRP4

PGE_1_ and PGE_2_ were found to stimulate MRP4 ATPase activity. Both are composed of 20 carbon atoms, organized as a cyclopentanone core (C8-C12) with two alkyl chain substituents (the α chain, C1-C7, and ω chain, C13-C20). These two PGs share a carboxyl at C1, a ketone at C9, hydroxyls at C11 and 15, and a carbon-carbon double bond at C13 (Fig. 1f). PGE_1_ differs from PGE_2_ only by the saturation of a second carbon-carbon double bond at C5, and both substrates share a similar pose when bound to MRP4. Though the map for PGE_2_-bound MRP4 is higher resolution than the PGE_1_-bound sample, either substrate can be confidently placed in its respective density. Both PGE_1_ and PGE_2_ bind MRP4 in the solvent-accessible central cavity, engaging residues from both TMD1 and TMD2. The binding site is a narrow channel composed of a large hydrophobic region and pockets of charged residues that coordinate the substrate’s polar groups (Fig. 5b).

**Fig 5.**
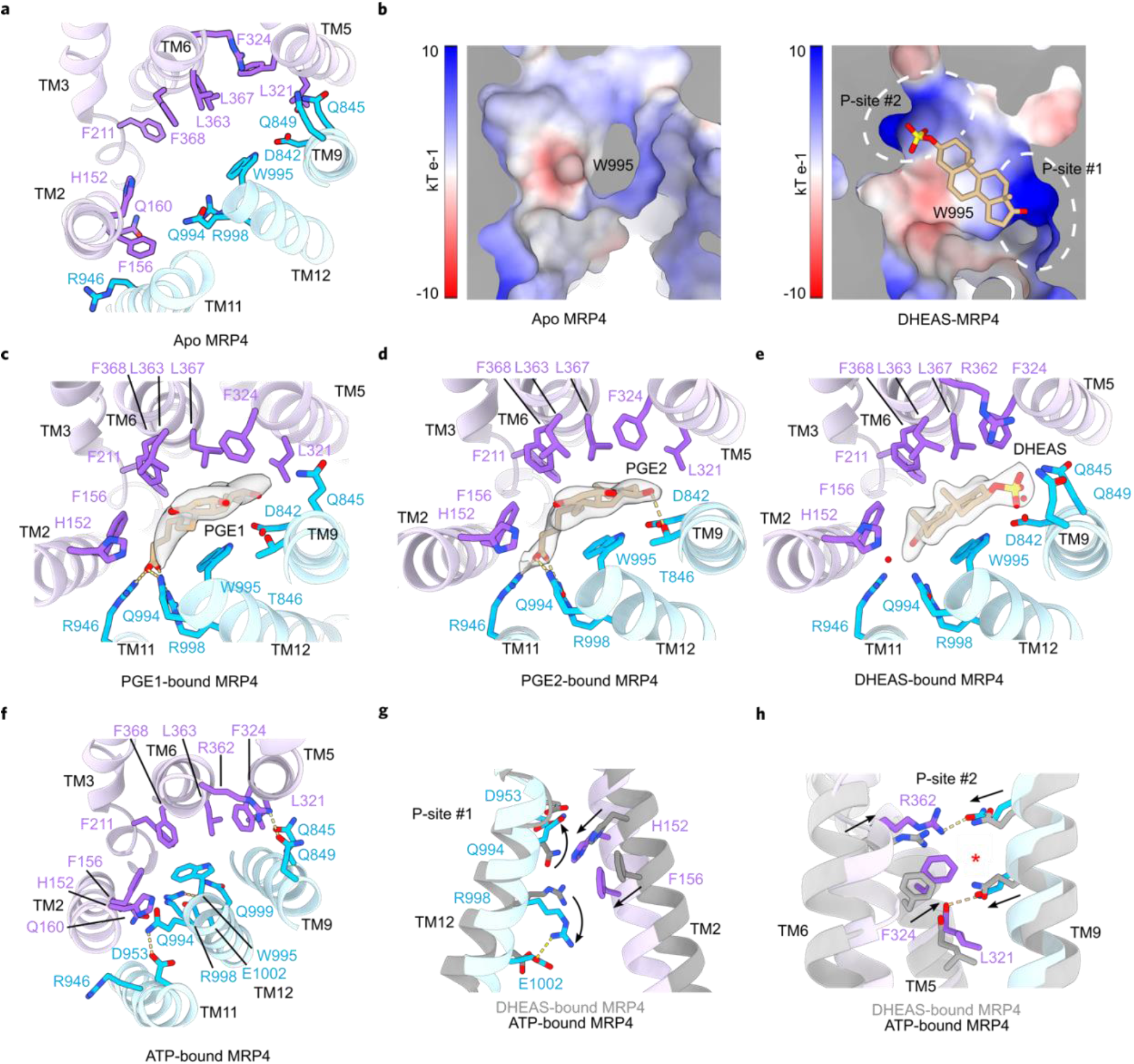
DHEA-S and prostaglandin binding to MRP4. Top-down view of the substrate binding pocket in **a**, the apo, inward-open state, **c**, the inward-narrow state bound to PGE_1_, **d**, the inward-narrow state bound to PGE_2_, **e**, the inward-narrow state bound to DHEA-S, and **f**, the outward-occluded state of MRP4_E1202Q_ bound to ATP-Mg^2+^. Residues shown form van der Waals contacts, hydrogen bonds, or salt bridges with any of the three substrates. Substrates are represented by sticks. For clarity, only substrate densities are shown as transparent surfaces. Waters are depicted as red spheres, and electrostatic interactions are indicated as yellow dashed lines. **b**, Electrostatic potentials of the substrate-binding surfaces of MRP4’s TMD2 in the apo, inward open state (left) and the inward-narrow, DHEA-S-bound state (right). Dashed circles highlight P-sites #1 and #2 that assemble upon substrate binding. Local conformational changes at **g**, P-site #1 and **h**, P-site #2 as MRP4 transitions from the inward-narrow, substrate-bound state to the outward-occluded, ATP-Mg^2+^-bound state. The DHEA-S bound structure is shown in gray and the ATP-Mg^2+^-bound state is colored as in Fig. 1a. Black arrows indicate movement of sidechains. Electrostatic interactions are shown as yellow dashed lines. Red asterisks indicate positions of substrate polar groups in their respective structures (an unoccupied P-site #2 and the C1 carboxyl in P-site #1 in the PGE1 and PGE2 bound structures; the 3β sulfate in P-site #2 and a coordinated water in P-site #1 in the DHEA-S bound structure).

In either PG-bound structure, the alkyl chains and the cyclopentanone core are wedged between W995 on TMD2 and a hydrophobic patch on TMD1. F156, F211, F324, L363, L367, and F368 form extensive van der Waals contacts with the 20-carbon backbone. The shared polar substituents of PGE_1_ and PGE_2_ serve to further orient the substrates. The C1 carboxyl sits in a positively charged pocket (P-site #1) located between TM 2, 11, and 12. The group forms salt bridges with R946 and R998, and maintains additional electrostatic interactions with H152, Q160 and Q994. The C11 hydroxyl and the C10 ketone substituents of the cyclopentanone ring point toward a narrow solvent-accessible cavity that extends above the substrates toward the extracellular half of MRP4, while the C15 hydroxyl of either PG faces a pocket on TM9 formed by D842, Q845, and T856. Taken together, the combination of a hydrophobic core, the α chain carboxyl, and the C10, 11 and 15 substituents allows MRP4 to bind and translocate these PGs specifically.

In addition to these two prostanoids, we also found that DHEA-S stimulated MRP4 ATPase activity. The sterol core of DHEA-S is conjugated by a sulfate ester at the 3β position and has a ketone substituent on C17, providing polar groups at either end of the aliphatic sterol core for electrostatic interactions. The map of DHEA-S-bound MRP4 is of high resolution (2.7 Å), allowing for definitive assignment of the substrate and relevant sidechain conformations. MRP4 binds DHEA-S in the same site as PGE_1_ and PGE_2_, using residues from both TMD1 and TMD2. The binding site residues in all three substrate-bound structures adopt similar rotamers, revealing a common substrate-bound state.

The 17-ketosteroid backbone of DHEA-S makes extensive van der Waals contacts through its β face with the hydrophobic patch formed by residues from TMD1. Residues L363, L367, and F368 intercalate between the two methyl substituents of DHEA-S, while F156, F211, and F324 pack against the periphery of the substrate’s sterol core. W995 from TM12 stacks against the α face of DHEA-S, and the indole ring of W995 hydrogen bonds with the carboxyl of D842. The basic P-site #1, composed of H152 from TM2, R946 from TM11, and R998 and Q994 from TM12, surrounds the steroid’s ketone moiety and an ordered water. The 3β sulfate lies in a distinct pocket of positive charge (P-site #2) at the interface of TM6 and TM9, where R362 and Q849 form electrostatic interactions with one of the sulfate’s three non-bridging oxygens. A second oxygen is coordinated by Q845 and makes a water-mediated interaction with D842, creating a network of interactions that links the sulfo group to W995. The third oxygen is not coordinated by the protein but instead faces a narrow solvent-filled tunnel that continues in the direction of the extracellular face. Neither PGE_1_ nor PGE_2_ make notable contacts with the sidechains directly involved in binding the sulfo-group of DHEA-S.

Although the two helical bundles clamp around the substrates in all three structures, there is little change to the rotamers of most substrate-binding residues compared to the apo state. The few residues that adopt new conformations include F156 and F324, which flip to line the periphery of the hydrophobic face of TMD1. The repositioning of F156 allows for R946 to rotate from facing out of the helical bundle (toward the lasso motif, partially resolved in the DHEA-S and PGE_1_ bound structures) to facing in toward the central cavity, where its guanidino group contributes to P-site #1. A similar inward rotation by H153 adds further positive-charge character to the pocket relative to the apo state (Fig. 5b).

In the ATP-bound state, MRP4’s transmembrane regions are in an arrangement incompatible with substrate binding. The two halves of the transporter have rotated inward toward the central cavity and F211, L367, and F368 from the hydrophobic face of TMD1 are within van der Waals contact distance of W995, sterically preventing DHEA-S or either PG from occupying the site (Fig. 5f). ATP-binding also results in the intrusion of TMD2 hydrophobic residues into the regions of positive charge at either end of the substrate binding pocket. P-site #1 is disrupted by the insertion of F156 and H152, displacing the substrate binding residues Q994 and R998. Q994 rotates away to neutralize D953 from TM11, while R998 adopts electrostatic interactions with Q160 and Q999 and forms a salt bridge with E1002 (Fig. 5g). At the other end of the substrate binding pocket, F324 occupies the former site of the DHEA-S sulfo group in P-site #2 (Fig. 5h). Of the residues previously involved in coordinating the sulfate’s oxygens, R362 and N849 hydrogen bond with one another, while N845 flips away and is stabilized by the backbone carbonyl of L321. The tight packing observed at both P-sites and between the hydrophobic face on TMD1 and W995 from TMD2 is consistent with substrate expulsion from the binding site, though no clear exit pathway is present

### Structural comparison between MRP4 and the leukotriene transporter MRP1

MRP4 shares 37% sequence identity with the closely related and well-studied MRP1 (Supplementary Fig. 1). While both are recognized as transporters of amphipathic organic anions, only MRP1 is well described as a transporter of glutathione conjugates, including the eicosanoid LTC_4_. In our studies, LTC_4_ failed to stimulate MRP4, unlike reported effects on MRP1^45^. Differences in the binding sites of substrate-bound MRP4 and MRP1 are consistent with their respective substrate specificities. The structure of DHEAS-bound MRP4, when superimposed onto LTC_4_-bound MRP1 (PDB ID: 5UJA), aligned via TM bundle 2, shows that the MRP4 substrate binding site is narrower and lacks shape complementarity to the leukotriene (Fig. 6a). TM bundle 1 of MRP4 is rotated toward the central axis of the transporter by ~13° compared to its position in MRP1, resulting in an apparent clash between TM2 residues in MRP4 and the glutathione moiety of LTC_4_ (Fig. 6b).

**Fig 6.**
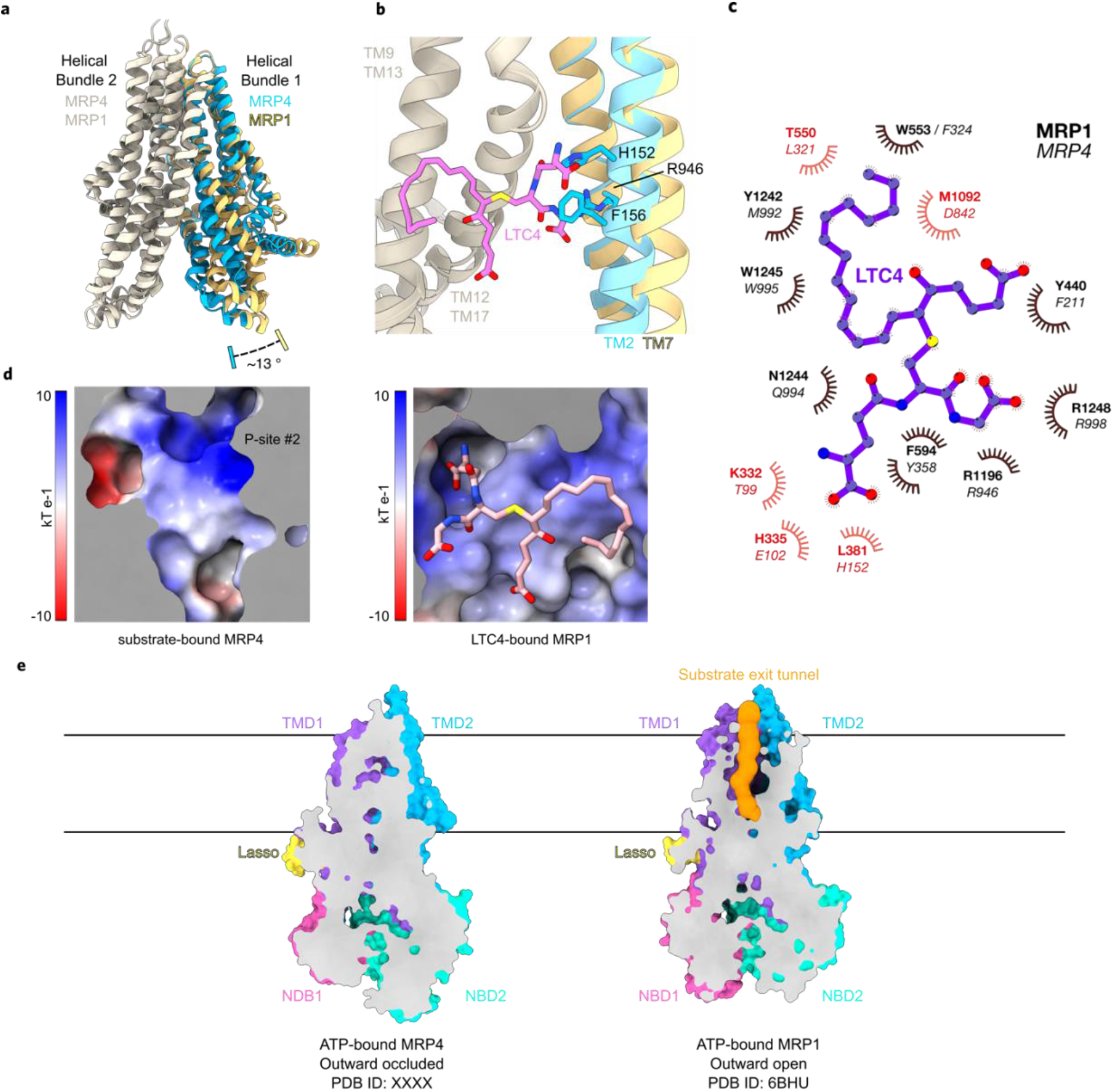
Structural comparison between MRP4 and related ABCC family members. **a**, Transmembrane domains of MRP1 and MRP4 aligned through helical bundle #2. Helical bundle #2 is colored bone for both transporters. Helical bundle #1 is colored canary in MRP1 and cyan in MRP4. The relative angle of rotation between MRP1’s and MRP4’s helical bundle 1 with respect to helical bundle 2 is indicated. **b**, Close-up view of the superimposed structures of LTC4-bound MRP1 and DHEAS-bound MRP4 (DHEAS not shown for clarity). LTC4 sterically clashes against H152, F156, and R946 as a result of MRP4’s narrower substrate-bound conformation. Transmembrane regions colored as in **a**, LTC4 and MRP4’s sidechains shown as sticks. **c**, Schematic of MRP1 bound to LTC4 (PDB:5UJA). MRP1 residues involved in ligand binding are labeled in bold, while aligned MRP4 residues are noted in italics. Red text signified a change in polarity between MRP1 and corresponding MRP4 residues. **d**, Electrostatic potentials of the aligned substrate-binding surfaces of MRP4’s TMD2 bound to DHEA-S (not shown for clarity) (right) and MRP1’s TMD2 bound to LTC4. An acidic patch in MRP4 replaces a basic surface for coordinating glutathione carboxyl groups. LTC4 shown as sticks, colored magenta and by heteroatom. **e**, Slice-through of ATP-bound MRP4 and ATP-bound MRP1, reveals a substrate exit tunnel present only in the MRP1 structure, while MRP4 is outward-facing occluded. Transmembrane regions of MRP4 and aligned regions of MRP1 are colored as in Figure 1. TMD0 of MRP1 hidden for clarity. The volume of the MRP1 substrate exit tunnel is shown as an orange surface.

Per residue differences between the binding pockets of MRP4 and MRP1 may further explain the lack of LTC_4_ stimulation by MRP4. Of the fifteen MRP1 residues involved in LTC_4_ binding, ten are identical or similar at the equivalent positions in MRP4, while the remaining five are altered (Fig. 6c). Two MRP1 residues (K332, H335) that hydrogen bond with a carboxyl from the LTC_4_ glutathione moiety are replaced by a threonine and a glutamic acid in MRP4, forming an acidic pocket in place of the basic pocket in MRP1 (Fig. 6d). M1092, part of the hydrophobic region of the MRP1 binding site that packs against the lipid tail of LTC_4_, is equivalent to D842 in MRP4, which has a crucial role in orienting polar groups of DHEA-S and the PGs. T550 in MRP1 is replaced by L321 in MRP4, which could sterically clash with LTC_4_ in the binding site. Together, these differences can account for the lack of substrate specificity of MRP4 for glutathione conjugates while maintaining transport of other organic anions.

The molecular structure of ATP-bound MRP4 illuminates a new state along the ABCC family substrate transport cycle, differing substantially from the ATP-bound state of MRP1 (PDB ID: 6BHU) (Fig. 6e). Whereas MRP1 is outward open upon ATP-binding and contains a narrow channel extending between helices 6,11,12 and 17 from the substrate binding residues to the extracellular milieu, the ATP-bound MRP4 structure contains a closed extracellular gate with no apparent substrate exit tunnel. The closure of this extracellular pathway is independent of ATP hydrolysis or subsequent phosphate release and supports the ‘alternating-access’ mechanism for directionality of substrate transport where the extracellular face of the exporter resets prior to the cytoplasmic portions of the transporter.

## DISCUSSION

Our series of structures offer insights into the mechanism of substrate recognition and translocation by the organic anion transporter MRP4 (Fig. 7a). Based on our structures and on the closely related MRP1_E1454Q_ bound to ATP-Mg^2+^, we propose the following translocation model for MRP4. In the inward-open apo state, the amphipathic substrate-binding site is located in the large central cavity between two helical bundles. The hydrophobic region on TMD1 and at least two distinct pockets of positive charge allow MRP4 to recognize organic anions of varying chemical scaffolds. In our PG and DHEA-S bound structures, residues from both TMDs engage the substrates, bringing the two helical bundles in toward the central axis of the transporter. The inter-NBD distance is shortened by up to 19 Å in the resulting inward-open narrow conformation, priming the NBDs for an accelerated rate of dimerization in the presence of ATP-Mg^2+^. Nucleotide binding drives the global conformational transition that results in substrate transport across the membrane. Our ATP-bound MRP4_E1202Q_ features dimerized NBDs and no observable exit pathway in the TMD region linking substrate-binding residues to the extracellular space. Consistent with an alternating access model, this structure represents a state subsequent to substrate translocation, where the extracellular gate has closed and ATP-hydrolysis and phosphate release have not yet reset the cytoplasmic portions of the transporter.

**Fig 7.**
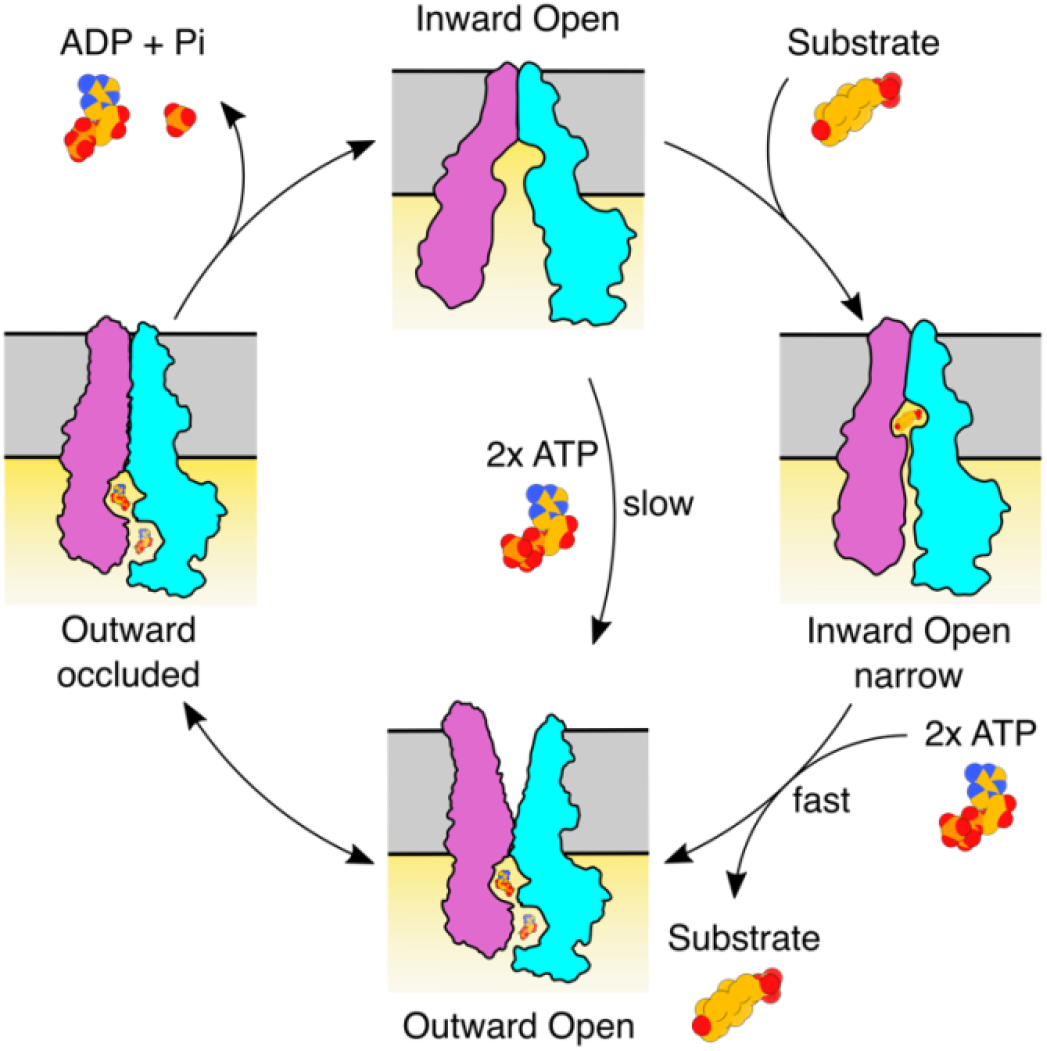
Proposed translocation cycle of MRP4. Cartoon representation of the alternate access transport mechanism of MRP4, derived from current cryo-EM structures and the ATP-bound MRP1 structure (PDB ID: 6BHU). The cycle begins with MRP4 open toward the cytoplasm, featuring a solvent-accessible substrate binding cavity and widely separated NBDs. When bound to substrate, the two halves of MRP4 constrict toward one another, reducing the volume of the central cavity and bringing the NBDs into close proximity. ATP binding dimerizes the NBDs and induces substrate transport by allosterically deforming the substrate binding-site and opening the extracellular gate for substrate release. Inward open MRP4 is capable of binding ATP in the absence of substrate and proceeding to this outward open conformation, though at notably slower rate due to the separation in NBDs. In the outward occluded state, thermodynamic fluctuations reset the exit pathway, leaving the substrate binding site exposed to neither the cytoplasm nor extracellular space. ATP hydrolysis and subsequent nucleotide and Pi release allows for dissociation of the NBD dimer and return to the inward open state. The first half of MRP4 composed of the L0, TM bundle 1, and NBD1 shown in magenta, and the second half of MRP4 composed of TM bundle 2 and NBD2 shown in cyan.

Our ATP-bound MRP4 structure stands in contrast to that of ATP-bound MRP1, where cryo-EM analysis of the analogous Walker A mutant MRP1_E1454Q_ in the presence of ATP-Mg^2+^ exclusively resolved an outward-open transporter. Extensive processing of our MRP4_E1202Q_ dataset provided no evidence for an outward-open conformation. Several 2D classes, which together represent a minority of total particles, show MRP4 to be inward-open, and the low-resolution map using particles from those classes matches our apo structure (Supplementary Fig. 7a). These results suggest that the outward open conformation of MRP4 may be more transient than that of MRP1, representing subtle differences in the energy landscape of the two related transporters.

Outward-occluded states have been reported for the type IV bacterial exporters MsbA^46^, TmrAB^47^, Cgt^48^, and McjD^49^, as well as the eukaryotic phosphatidylcholine exporter ABCB4^50^. Though the precise helical organization differs between each transporter, these outward-occluded structures are thought to represent similar states in the above model of substrate transport. The structure of ATP-bound MRP4 most strongly resembles the conformation of the ADP/ATP bound state of the ABCC family member SUR1 (Supplementary Fig. 10a-c), a sulphonylurea receptor that acts as the ADP sensing subunit of the KATP channel^51^. Despite SUR1 sequence homology to the MRP subfamily and overall structural resemblance to a canonical ABC transporter, it has no known transport substrates and is thought to exclusively regulate channel activity. The ATP-bound MRP4 and ADP/ATP-bound SUR1 both contain a closed extracellular gate and tightly packed TMDs sealed off from the cytoplasm, suggesting conformational similarity of MRP4 to the ABCC family extends beyond the MRPs.

We show that PGE_1_, PGE_2_ and DHEA-S are preferred over other previously reported substrates, stimulating MRP4 ATPase activity several fold with apparent K_m_ values in the micromolar range. Our substrate-bound structures are consistent with this high affinity, revealing a binding site that can accommodate multiple species of organic anions. MRP4 sequesters the non-polar cores of both PGs and DHEA-S between the hydrophobic regions of TMD1 and W995 on the opposing face of the pocket. This interaction mimics the ‘tryptophan-sandwich’ observed in the LTC4-MRP1 bound structure^45^, and is likely a common feature across the MRP subfamily used to bury hydrophobic cargo. The substrate-induced transition in MRP4 from the inward open apo- to inward open-narrow results in the formation of two distinct positively charged pockets, either through sidechain flipping (as in P-site #1 with R946) or a simple clamping of the TM bundles (as in P-site #2). Both P-sites are capable of coordinating anionic groups, and their positioning at opposite ends of the binding site likely contributes to the diversity of substrates recognized by MRP4. All three substrate-bound structures are consistent with previous MRP4 mutagenesis studies that identified critical roles for F368, W995 and R998^52,53^ in substrate transport.

Our two PG-bound structures reveal comparable interactions between MRP4 and the C1 carboxyl, cyclopentanone core, and the C15 hydroxyl of either PG. Given that these functional groups are common to the other prostanoids shown to be transported by MRP4, including PGD_2_, PGF_2α_, and TXB_2_^17,19,20^, it is likely that they bind through similar contacts. Furthermore, these chemical features are shared by many other PGs, suggesting the diversity of PGs and their analogs transported by MRP4 may be currently underappreciated.

The result that PG and DHEA-S stimulate MRP4 ATPase activity contrasts with the remaining substrates tested, including the cyclic nucleotides. Despite reports which identify MRP4 as a nucleotide transporter^24,54,55^, high concentrations of cAMP and cGMP did not notably stimulate MRP4 ATPase activity in vitro. Our cryoEM results failed to confirm a high affinity complex between cAMP and MRP4, with the final reconstruction lacking both an observed density for cAMP and any notable conformational change compared to the inward-open apo state. While we cannot rule out cAMP as an MRP4 substrate, these results suggest that any transport of cyclic nucleotides in our reconstituted system occurs at or below the basal activity of MRP4, and that cAMP binding does not induce an inward-open narrow state. Downstream signaling of various PGs through their respective GPCRs affects the levels of cytoplasmic cAMP^56^, providing one possible mechanism for regulation of this cyclic nucleotide by MRP4. Further investigations will be necessary to clarify the precise molecular details of transport of cAMP and other non-stimulating substrates by MRP4.

Our study provides the first atomic-level descriptions of MRP4 in three distinct conformations and reveals the basis of its organic anion specificity. The structures of PGE_1_ and PGE_2_ bound to MRP4 underscore the role of this transporter in prostaglandin efflux, a signaling mechanism central to numerous physiological processes. The OF-occluded structure fills in a previously unobserved but assumed state of the MRP translocation cycle and can be used as the basis for homology models of other MRPs. Our series of IF and IF-narrow structures will provide the groundwork for computational studies aimed at validating endogenous and xenobiotic substrates of MRP4 and will aid in structure-guided designs of inhibitors for this promiscuous transporter. Improved understanding of MRP4 function will aid in addressing multidrug resistance and transporter dysregulation.

## METHODS

### MRP4 expression and membrane preparation

The wild-type bovine MRP4 gene and the MRP4_E1202Q_ mutant were synthesized by Genscript and cloned into pFastBac with a C-terminal thrombin-cleavable 8xHis tag. For all constructs, recombinant baculovirus was produced using the Bac-to-Bac Baculovirus Expression System (Thermo Fisher Scientific). High-titer (>10^9^ viral particles/mL) P3 virus was used to infect *Spodoptera frugiperda (Sf9)* cells at a cell density of 3×10^6^ cells mL^-1^. Cells were harvested 48 hours post infection by centrifugation at 2000 *g*, flash frozen and stored at −80°C until use. Frozen cell pellets were thawed and resuspended in lysis buffer, 50 mM Tris (pH 8.0), 300 mM NaCl, 1 mM TCEP, 1 mM phenylmethylsulfonyl fluoride (PMSF), and EDTA-free cOmplete protease inhibitor cocktail tablets (Sigma-Aldrich). The resuspended cells were repeatedly homogenized with a Dounce homogenizer, then sonicated on ice at 1 s^-1^ for 5 minutes. Membranes were spun down at 100,000 *g* for 90 minutes, then resuspended in working buffer (50 mM Tris (pH 8.0), 200 mM NaCl, 1 mM TCEP), supplemented with 1 mM PMSF, with EDTA-free cOmplete protease inhibitor cocktail, and flash frozen in aliquots for future use.

### MRP4 nanodisc reconstitution

Thawed membranes were solubilized with 1% (w/v) *n*-dodecyl-*β*-d-maltopyranoside (DDM, Anatrace) added at a detergent to protein ratio of 0.2 (w/w) and stirred for 4 h. The supernatant was separated from insoluble fractions by centrifugation at 100,000 *g* for 30 minutes and clarified through a 0.22 μm filter. After adding 5 mM imidazole, the supernatant was batch bound to TALON IMAC resin (Clontech) for 2 h at 4°C. The resin was washed with 10 column volumes of working buffer supplemented with 10 mM imidazole and 0.05% DDM, and then returned to working buffer supplemented with 5 mM imidazole and 0.05% DDM. Resuspended resin with bound MRP4 was mixed with Soybean Polar Lipid Extract (Avanti) to a final estimated lipid concentration of 1.2 mM and mixed for 1 h at room temperature. Purified MSPE3D1 was added to the mixture at a final concentration of 0.6 mg ml^-1^ and mixed for 1 h at room temperature. Finally, 60 mg ml^-1^ of methanol-activated Bio-Beads SM2 resin (Bio-Rad) was added and allowed to mix overnight at 4°C. The heterogenous resin mixture was washed with resuspension buffer extensively to remove empty nanodiscs, and resin-bound MRP4-nanodisc complexes were eluted using buffer supplemented with 150 mM imidazole. The elution fraction was concentrated at 2000 *g* in a 100 kDa Amicon Ultra concentrator (Millipore), filtered, and purified by SEC on a Superdex 200 Increase column (Cytiva) in 20 mM Tris pH 8.0, 150 mM NaCl, 1 mM TCEP. All steps were performed at 4°C.

### ATPase Assay

ATP hydrolysis by MRP4 was observed using an established NADH-coupled ATPase assay^57^ by monitoring λ_ex_ = 340 nm and λ_em_ = 445 nm on a SpectroMax plate reader. MRP4-nanodisc (400 nM), 60 mg mL^-1^ pyruvate kinase (Sigma-Aldrich), 32 mg mL^-1^ lactate dehydrogenase (Sigma-Aldrich), 4 mM phosphoenolpyruvate, and 150 mM NADH were mixed in 50 mM Tris pH 8.0, 150 mM KCl, 2 mM MgCl_2_ buffer. For the ATP concentration dependency studies, ATP was added at the specified concentrations. For investigation of substrate concentration dependence, ATP was held constant at 4 mM while substrate was added at the specified concentrations. The resulting data were analyzed using GraphPad Prism to fit the Michaelis-Menten equation.

### Cryo-EM sample preparation and data acquisition

Freshly purified MRP4 in lipid nanodiscs was concentrated to 0.6 mg ml^-1^ for vitrification. Quantifoil R1.2/1.3 400-mesh Cu Holey Carbon Grids (EMS) were glow discharged at 15 mA for 30 s immediately before use. Sample was applied and grids were blotted using a Mark IV Vitrobot (Thermo Fisher Scientific) for 5 s at 100% humidity and 4°C before plunge freezing in liquid N_2_-cooled ethane. For the ATP-bound state, MRP4_E1202Q_ in lipid nanodiscs was treated similarly, except for the addition of 1 mM PGE_2_ and 10 mM ATP, 10 mM MgCl_2_ followed by a 10 min incubation at 37°C prior to applying on grids. The substrate-bound samples were prepared similar to apo, except for the use of MRP4_E1202Q_ in lipid nanodisc and the inclusion of 200 μM of either DHEA-S, PGE_1_, or PGE_2_ or 400 μM cAMP. All three substrate-bound samples were blotted on Quantifoil R1.2/1.3 300-mesh Au Holey Carbon Grids (EMS). For all samples, grids were screened for ice quality and particle density using a Talos Arctica (Thermo Fisher Scientific) microscope.

For the nucleotide-free MRP4-nanodisc sample, 4698 118-frame super-resolution movies were collected on an FEI Titan Krios (Thermo Fisher Scientific) operated at 300 keV, equipped with a Bioquantum energy filter (Gatan) set to a slit width of 20 eV and a post-GIF K3 camera in single-electron counting mode. The dataset was collected with a beam image-shift over a 3×3 hole array at a nominal magnification of 105,000x and physical pixel size of 0.834 Å pix^-1^, using an underfocus range of 1.0 to 2.0 μm. SerialEM was used for all data acquisition using semi-automated scripts. The ATP-bound MRP4_E1202Q_ dataset was collected on a different FEI Titan Krios using similar parameters, except for a physical pixel size of 0.835 Å pix^-1^. That dataset was composed of 3176 super-resolution movies collected at an underfocus ranging from 0.5 to 2.0 μm.

All substrate-bound data was collected on this second FEI Titan Krios after an upgrade to Fringe-Free Imaging using a beam image-shift over a 3×3 hole array and recording 3 movies per hole. All datasets used an underfocus range of 0.5 to 2 μm. The nominal magnification and physical pixel size after upgrade were unchanged. For the DHEA-S-bound MRP4 sample, 5609 80-frame super-resolution movies were collected. This data was merged with a dataset of 1266 movies collected at 35° tilt, collected using a single record per hole. 7219 80-frame super-resolution movies were collected on the PGE_1_-bound MRP4 sample and merged with 1034 80-frame super-resolution movies collected at 35° tilt collected using a single record per hole. Finally, 8649 80-frame super-resolution movies were collected on the PGE_2_-bound MRP4 sample.

### Image processing

MotionCor2^58^ was used to correct all movie stacks for beam-induced motion, to sum frames with and without dose weighting, and to Fourier bin images 2 × 2 to the counting pixel size. The contrast transfer function (CTF) and resolution estimates for each corrected and dose weighted micrograph was determined in cryoSPARC^59^ using PatchCTF. Particles were picked in cryoSPARC 3.2 using a Gaussian disk as a template, and subsequently processed in cryoSPARC 3.2 as depicted in workflows shown in Supplementary Fig. 3-7.

### Model building and structure refinement

All five atomic models were built using Coot^60^, PHENIX^61^ and ISOLDE^62^. For the structures of apo-MRP4 and ATP-bound MRP4_E1202Q_, transmembrane helices and linkers were manually built in Coot using the output densities from DeepEMhancer^63^. The NBD models of the closely related bovine MRP1 (PDB: 6BHU), with residues mutated to those of MRP4 via Modeller^64^, were initially used to build the NBDs of ATP-bound MRP4_E1202Q_. The final ATP-bound MRP4 NBDs were used to guide model building of the NBDs of apo-MRP4 and all three substrate-bound structures, first by rigid-body fitting in ChimeraX, followed by real-space refinement in PHENIX and further correction in Coot and ISOLDE. All three substrate-bound states were built by rigid-body fitting individual domains of apo-MRP4 into the output densities from DeepEMhancer and following the steps as above for NBDs.

The deposited apo-MRP4 model contains residues 47-398, 409-419, 427-615, 692-745, 756-1298. The deposited PGE_1_-bound MRP4 model contains residues 23-394, 409-615, 693-745, and 756-1298, as well as a one PGE_1_ and one water molecule. The deposited PGE_2_-bound MRP4 model contains residues 46-394, 409-615, 693-745, and 756-1298, as well as one PGE_2_ and one water molecule. The deposited DHEA-S-bound MRP4 model contains residues 10-398, 409-615, 693-745, and 756-1298, as well as one DHEA-S and three water molecules. The deposited nucleotide-bound MRP4_E1202Q_ model contains residues 48-394, 408-615, 696-746, 756-882, and 898-1298 as well as two bound ATP, two Mg^2+^ ions, and a phosphatidylethanolamine lipid molecule. All models were validated against the sharpened densities from cryoSPARC. Visualizations and figures were prepared using UCSF Chimera^65^, ChimeraX^66^ and Inkscape software. Cavity calculations were made with MOLE*online*^67^ and the 3V Server^68^. 2D diagrams of ligand-protein interactions were produced using LigPlot+^69^.

## Data availability

All five three-dimensional cryoEM density maps have been deposited to the Electron Microscopy Data Bank under accession numbers EMD-XXXX (apo MRP4), EMD-XXXX (PGE_1_-bound MRP4), EMD-XXXX (PGE_2_-bound MRP4), EMD-XXXX (DHEA-S-bound MRP4), and EMD-XXXXMRP4_E1202Q_). The coordinates for the atomic models have been deposited in the Protein Data Bank under accession numbers XXXX (apo MRP4), XXXX (PGE_1_-bound MRP4), XXXX (PGE_2_-bound MRP4), XXXX (DHEA-S-bound MRP4), and XXXX (ATP-bound MRP4_E1202Q_). See Supplementary Table 1 for more details. Source data provided with this paper.

## ACKNOWLEDGEMENTS

We thank Phuong Nguyen for assistance with virus production and Daniel Asarnow for helpful discussions regarding purification and reconstitution of MRP4. This work was supported by the National Institute of General Medical Sciences (5P01GM111126 and NIGMS GM24485 to RMS and R35GM140847 to YC). Ruchika Bajaj was supported in part by American Heart Association postdoctoral fellowship Award No. 19POST34370101. Mass spectrometry was performed in the UCSF Mass Spectrometry facility supported by NIH grant P41 GM103481. We thank David P. Bulkley, Glenn Gilbert, and Mathew Harrington for support with cryoEM data collection. All data was collected at the UCSF cryoEM facility, which is supported by NIH grants S10OD020054, S10OD021741 and S10OD026881.

## AUTHOR CONTRIBUTIONS

R.M.S., D.L.K., C.S.C, Y.C., and A.S. conceived this work; S.P., R.B. and G.M.K expressed and functionally characterized proteins; S.P., E.G. and M.G. collected and processed cyroEM data; S.P. and E.G. performed model building and refinement; S.P., R.B., E.G., R.M.S. and D.L.K. wrote the manuscript with contributions from all other authors.

