## Supplementary Information for "Structural Basis of Prostaglandin Efflux by MRP4"

| | **Apo MRP4** (EMD-XXXXX) (PDB XXXX) | **MRP4 + DHEAS** (EMD-XXXXX) (PDB XXXX) | **MRP4 + PGE1** (EMD-XXXXX) (PDB XXXX) | **MRP4 + PGE2** (EMD-XXXXX) (PDB XXXX) | **MRP4 + ATP-Mg$^{2+}$** (EMD-XXXXX) (PDB XXXX) |
|---|---|---|---|---|---|
| **Data collection and processing** | | | | | |
| Microscope | Titan Krios G3 | Titan Krios G3 | Titan Krios G3 | Titan Krios G3 | Titan Krios G3 |
| Imaging System | K3 BioQuantum, 20 eV slit | K3 BioQuantum, 20 eV slit | K3 BioQuantum, 20 eV slit | K3 BioQuantum, 20 eV slit | K3 BioQuantum, 20 eV slit |
| Voltage (kV) | 300 | 300 | 300 | 300 | 300 |
| Electron exposure (e–/Å$^2$) | 67 | 67 | 67 | 67 | 67 |
| Exposure time (s) | 5.9 | 5.9 | 5.9 | 5.9 | 5.9 |
| Underfocus range (μm) | -1.0 to -2.0 | -0.5 to -2.0 | -0.5 to -2.0 | -0.5 to -2.0 | -0.5 to -2.0 |
| Pixel size (super-resolution) | 0.4175 Å pix$^{-1}$ | 0.4175 Å pix$^{-1}$ | 0.4175 Å pix$^{-1}$ | 0.4175 Å pix$^{-1}$ | 0.4175 Å pix$^{-1}$ |
| Frames per movie | 117 | 117 | 117 | 117 | 117 |
| Movies collected | 4698 | 5609 | 7219 | 8649 | 3176 |
| Acquisition software | SerialEM | SerialEM | SerialEM | SerialEM | SerialEM |
| Symmetry imposed | C1 | C1 | C1 | C1 | C1 |
| Initial particle images (no.) | 1,448,148 | 5,330,431 | 7,912,699 | 3,910,205 | 840,205 |
| Final particle images (no.) | 660,807 | 360,511 | 122,978 | 187,047 | 104,051 |
| Map resolution (Å) | 3.1 | 2.7 | 3.5 | 2.9 | 3.1 |
| FSC threshold | 0.143 | 0.143 | 0.143 | 0.143 | 0.143 |
| **Refinement** | | | | | |
| Model resolution (Å) | | | | | |
| Masked | 3.2 | 2.9 | 3.6 | 3.1 | 3.2 |
| Unmasked | 3.3 | 3.0 | 3.8 | 3.2 | 3.3 |
| FSC threshold | 0.5 | 0.5 | 0.5 | 0.5 | 0.5 |
| Model composition | | | | | |
| No. Atoms (non-hydrogen) | 9109 | 9521 | 9418 | 9202 | 9155 |
| No. Residues | 1144 | 1188 | 1175 | 1152 | 1135 |
| No. Water | 0 | 3 | 1 | 1 | 0 |
| No. Ligands | 0 | XH0: 1 | XPG: 1 | P2E: 1 | MG: 2 ATP: 2 PTY: 1 |
| *B* factors (Å$^2$) min/max/mean | | | | | |
| Protein | 37.02/230.10/125.69 | 1.73/190.69/85.26 | 34.87/204.17/100.34 | 7.57/190.27/89.80 | 101.57/273.17/150.35 |
| Ligand | --- | 10.47/41.83/24.62 | 46.19/46.19/46.19 | 37.95/55.85/44.75 | 142.09/187.07/157.54 |
| Water | --- | 28.76/36.26/32.62 | 41.80/41.80/41.80 | 33.33/33.33/33.33 | --- |
| R.m.s. deviations | | | | | |
| Bond lengths (Å) (# > 4σ) | 0.006 (0) | 0.004 (0) | 0.005 (0) | 0.006 (1) | 0.005 (1) |
| Bond angles (°) (# > 4σ) | 0.687 (6) | 0.674 (0) | 0.831 (0) | 0.898 (3) | 0.876 (2) |
| **Validation** | | | | | |
| MolProbity score | 1.61 | 1.12 | 0.78 | 0.77 | 0.65 |
| Clashscore | 6.77 | 2.34 | 0.94 | 0.64 | 0.43 |
| Ramachandran plot | | | | | |
| Favored (%) | 96.47 | 97.46 | 98.03 | 97.73 | 98.67 |
| Allowed (%) | 3.53 | 2.54 | 1.97 | 2.27 | 1.33 |
| Outliers (%) | 0.00 | 0.00 | 0.00 | 0.00 | 0.00 |
| Rama Z-score (RMSD) | | | | | |
| Whole | (N=1134) -1.93 (0.22) | (N=1180) -1.48 (0.21) | (N=1167) -0.82 (0.21) | (N=1144) -0.13 (0.23) | (N=1125) -0.80 (0.22) |
| Helix | (N=759) -1.23 (0.17) | (N=764) -1.18 (0.16) | (N=767) -1.46 (0.15) | (N=744) -0.28 (0.17) | (N=745) -0.74 (0.16) |
| Sheet | (N=47) 0.82 (0.71) | (N=51) 0.38 (0.63) | (N=73) -0.49 (0.49) | (N=73) 0.59 (0.58) | (N=83) 0.35 (0.53) |
| Loop | (N=328) -1.27 (0.31) | (N=365) -0.26 (0.30) | (N=327) -0.40 (0.32) | (N=327) 0.72 (0.36) | (N=297) 0.37 (0.35) |

**Supplementary Table 1. | Cryo-EM data collection, refinement and validation statistics**

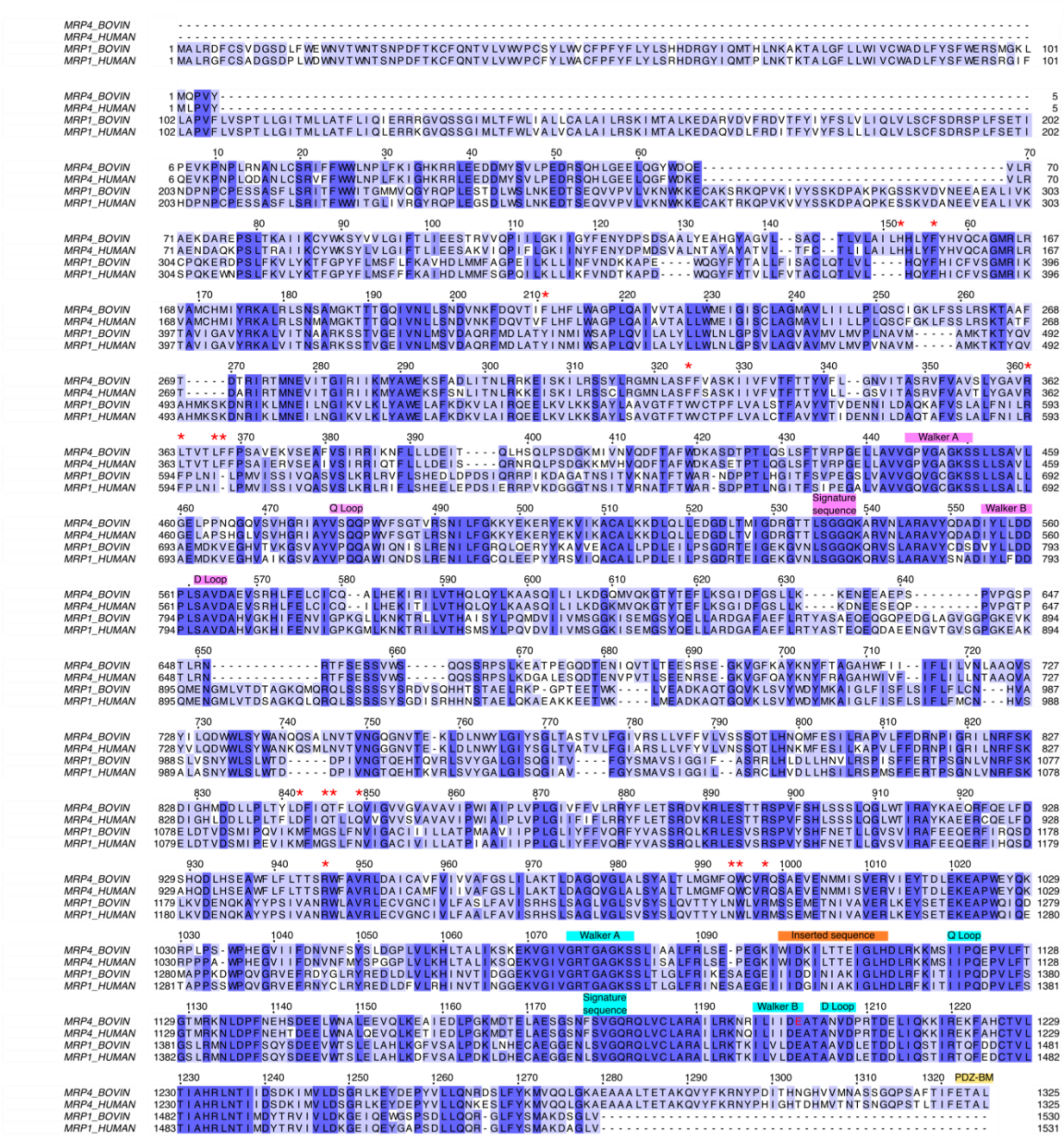

**Supplementary Fig 1. | Sequence alignment of MRP4 and MRP1 homologs**
Sequences of bovine and human MRP4 are aligned to reflect sequence conservation across evolution and similarity to the closely related MRP1. All references to MRP4 used throughout the text refer to the bovine sequence. The residue numbers for bovine MRP4 are indicated above the sequences. Red asterisks denote a residue involved in binding one of either DHEA-S, PGE1, or PGE2. Text boxes refer to conserved sequence motifs from NBD1 (pink) and NBD2 (cyan), NBD2's inserted sequence (orange) and the C-terminal PDZ-binding motif (PDZ-BM, sand). Bovine MRP4 catalytic residue E1202 is highlighted in red.

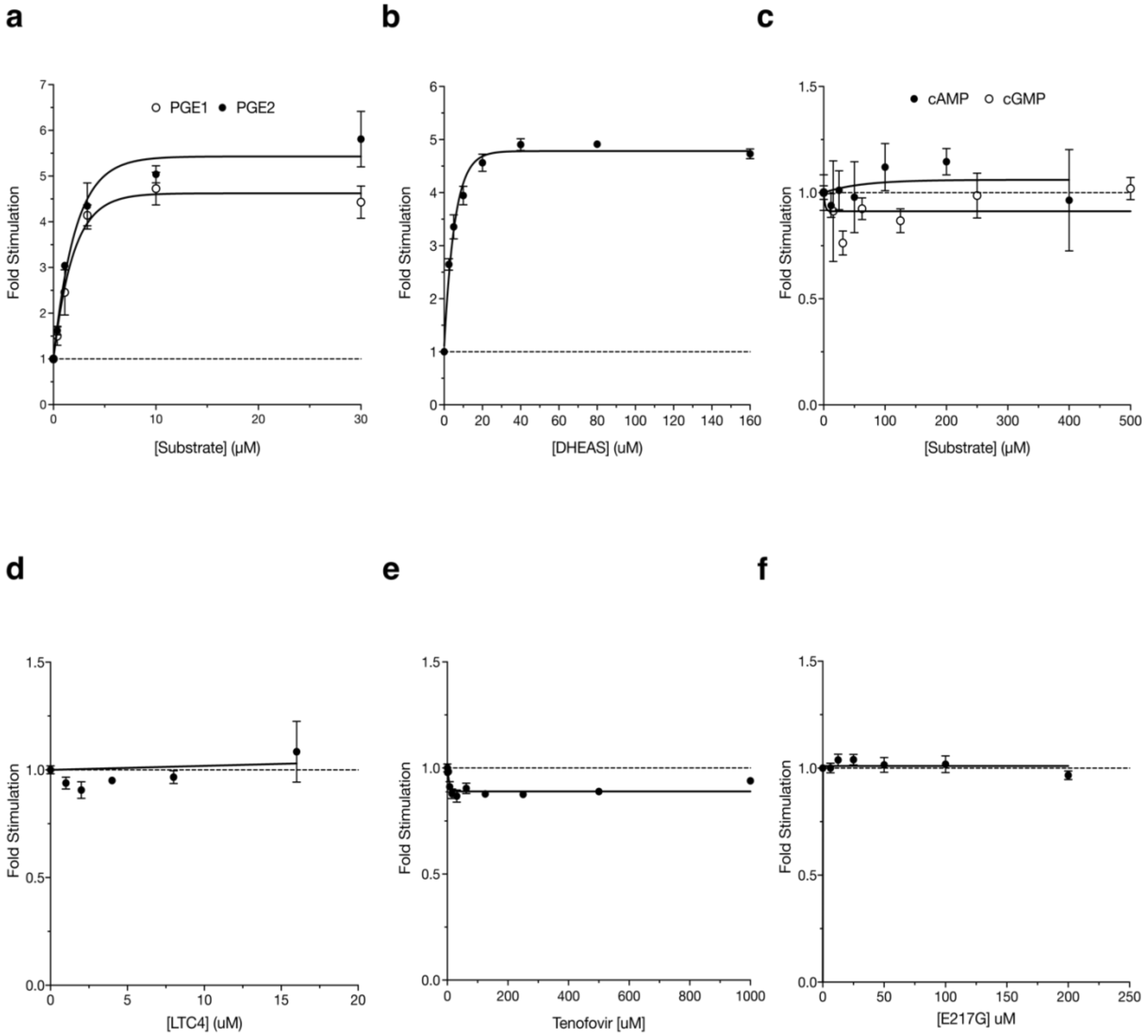


**Supplementary Figure 2. Biochemical characterization of MRP4.** Fold change in ATPase activity relative to basal levels with the addition of increasing concentrations of **a**, $PGE_1$, $PGE_2$, **b**, DHEAS, **c**, cAMP, cGMP, **d**, $LTC_4$, **e**, tenofovir and **f**, E217G. $PGE_1$, prostaglandin $E_1$; $PGE_2$, prostaglandin $E_2$; cAMP, cyclic adenosine monophosphate; cGMP, cyclic guanosine monophosphate; $LTC_4$, leukotriene $C_4$; DHEAS, dehydroepiandrosterone sulfate; E217G, β-estradiol-17β-D-glucuronide.

| Condition | $K_m$ (μM) (95% CI)[a] | $V_{max}$ (nmol $min^{-1}$ $mg^{-1}$) (95% CI)[a] | Maximal Fold Change |
|---|---|---|---|
| MRP4 + $PGE_1$ | 1.37 (0.84-2.28) | 57.76 (49.38-66.16) | 4.14 |
| MRP4 + $PGE_2$ | 1.68 (1.15-2.52) | 57.97 (52.27-63.70) | 5.05 |
| MRP4 + DHEAS | 2.32 (1.60-3.2) | 113.1 (106.6-119.9) | 4.78 |
| MRP4 + $LTC_4$ | NA | NA | 1.01 |
| MRP4 + E217G | NA | NA | 1.01 |
| MRP4 + cAMP | NA | NA | 1.07 |
| MRP4 + cGMP | NA | NA | 0.95 |

[a]ATPase activity in the presence of increasing concentrations of the indicated compounds was fit to a Michaelis-Menten equation. The kinetic parameters are mean values from at least three replicate experiments with 95% confidence intervals (CI). NA, not applicable.

**Supplementary Table 2: Kinetic parameters for ATPase activity of MRP4**

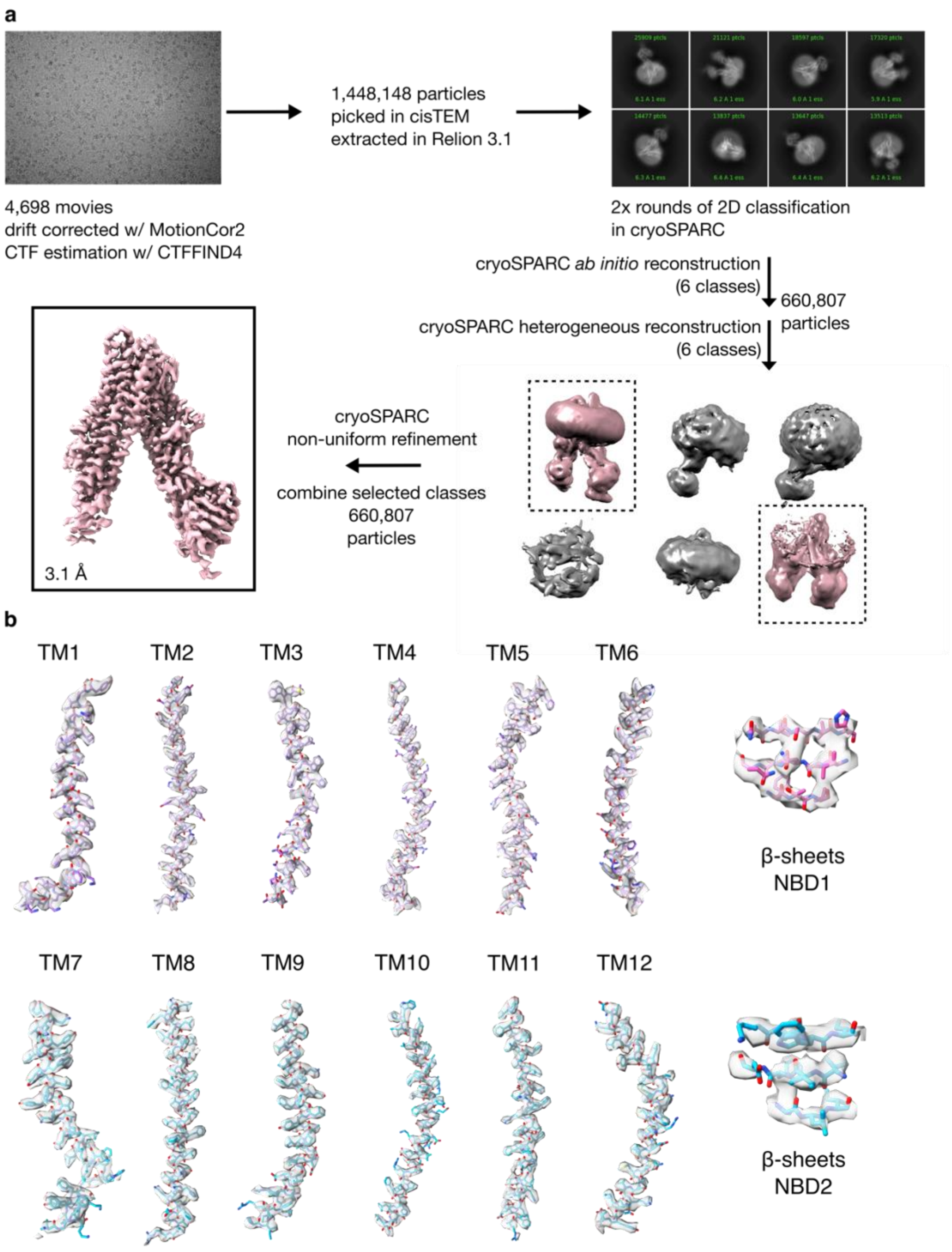
a
1,448,148 particles
picked in cisTEM
extracted in Relion 3.1
4,698 movies
drift corrected w/ MotionCor2
CTF estimation w/ CTFFIND4
2x rounds of 2D classification
in cryoSPARC
cryoSPARC *ab initio* reconstruction
(6 classes)
660,807
particles
cryoSPARC heterogeneous reconstruction
(6 classes)
cryoSPARC
non-uniform refinement
combine selected classes
660,807
particles
3.1 Å
b
TM1
TM2
TM3
TM4
TM5
TM6
β-sheets
NBD1
TM7
TM8
TM9
TM10
TM11
TM12
β-sheets
NBD2

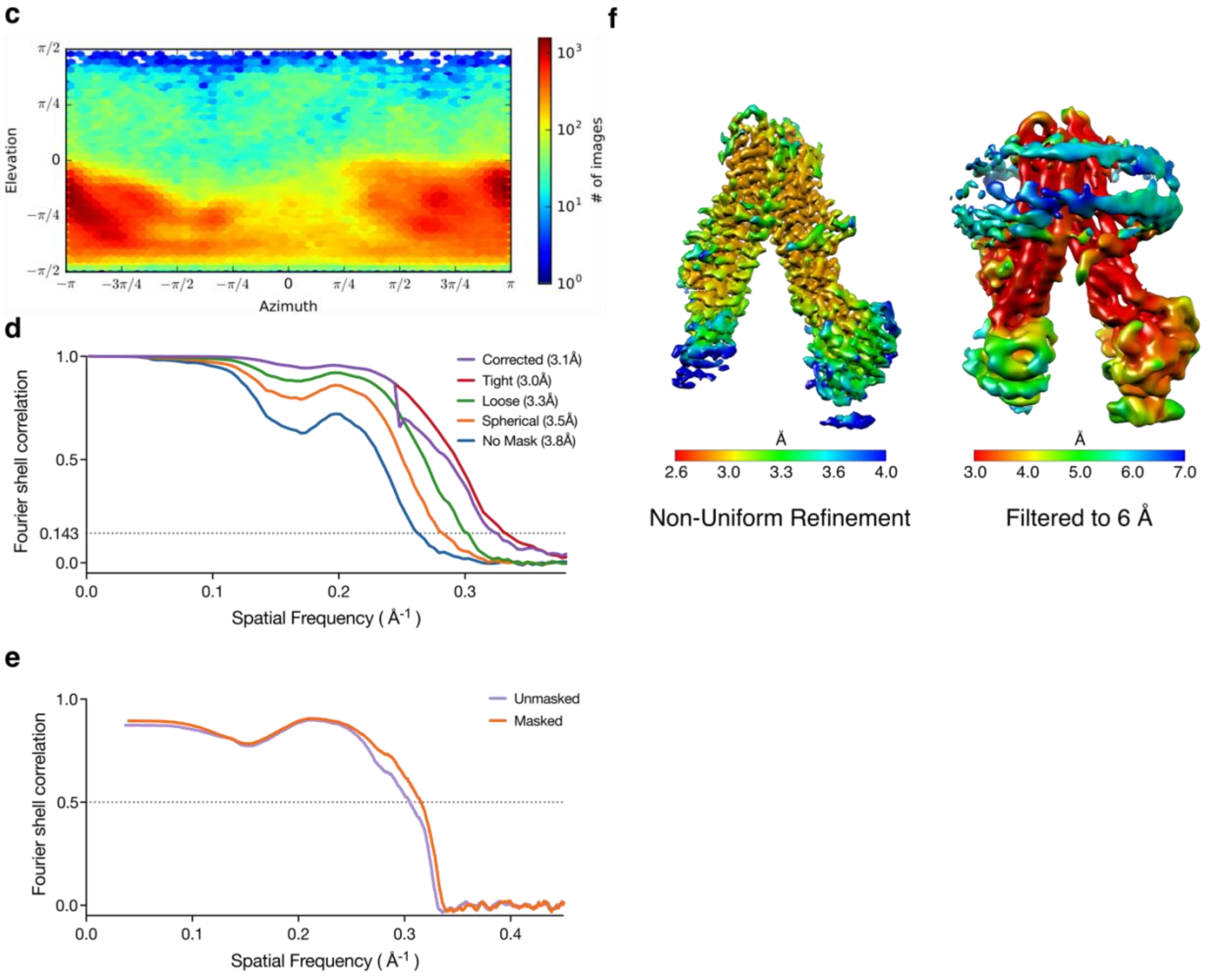


**Supplementary Figure | 3. Image processing of nucleotide-free, substrate-free MRP4 leading to a 3.1 Å cryo-EM reconstruction.** Cryo-EM analysis of apo MRP4 in lipid nanodiscs. **a**, Processing pipeline for apo MRP4. **b**, Cryo-EM density of the sharpened map overlaid with the model from selected regions. **c**, Euler angle heat-maps for the final nucleotide-free MRP4 refinement **d**, Gold-standard Fourier shell correlation threshold from the final nucleotide-free MRP4 refinement. Average resolution determined by using a correlation threshold of 0.143. **e**, Map to model FSC plot indicating correspondence of nucleotide-free MRP4 atomic model to final density map. **f**, Final density from cryoSPARC refinement colored by local resolution. To show lower resolution features, including NBD1, the map is also shown low-pass filtered to 6 Å.

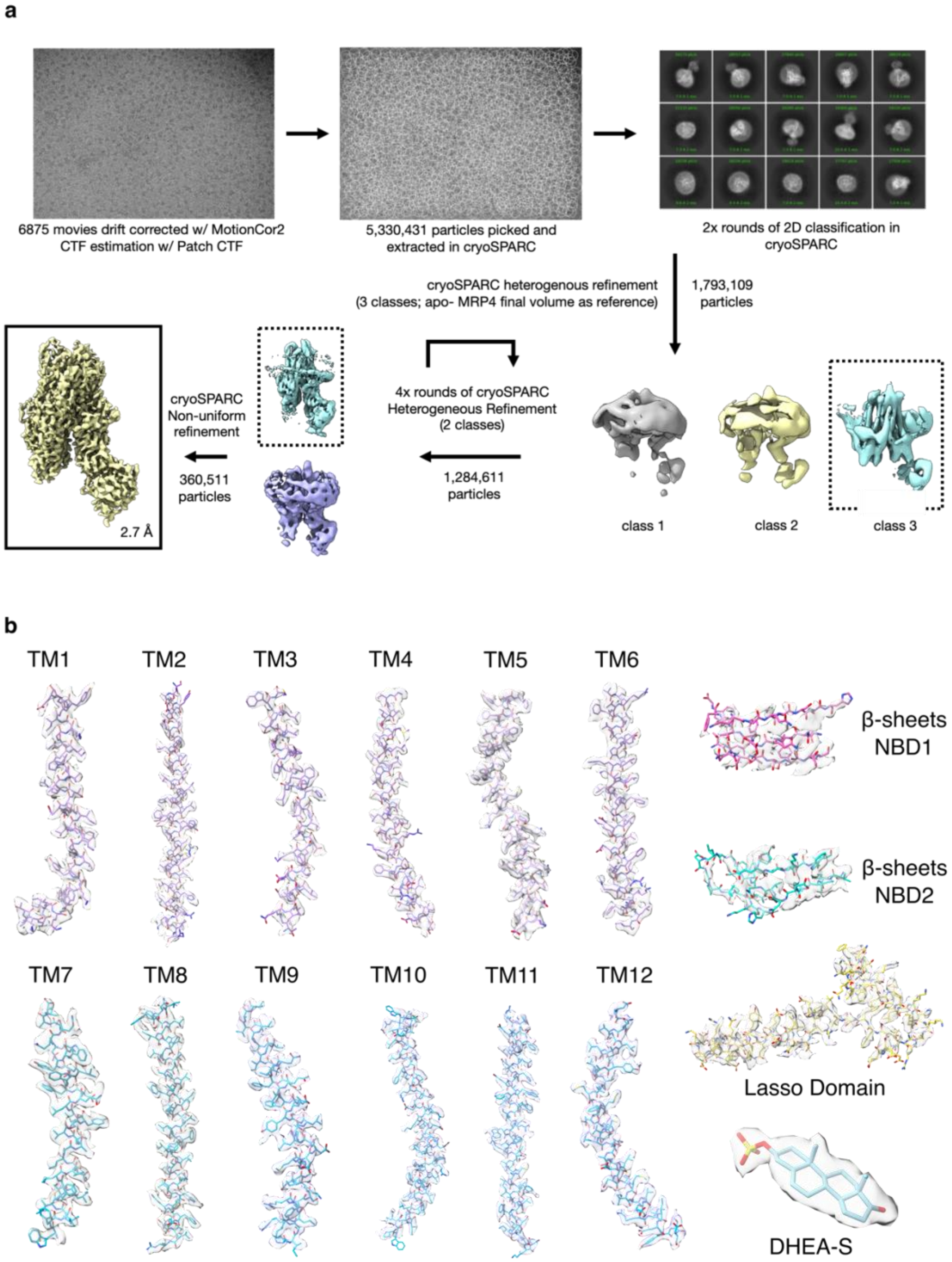
a
6875 movies drift corrected w/ MotionCor2
CTF estimation w/ Patch CTF
5,330,431 particles picked and extracted in cryoSPARC
2x rounds of 2D classification in cryoSPARC
cryoSPARC heterogenous refinement
(3 classes; apo- MRP4 final volume as reference)
1,793,109 particles
class 1
class 2
class 3
4x rounds of cryoSPARC Heterogeneous Refinement (2 classes)
1,284,611 particles
cryoSPARC Non-uniform refinement
360,511 particles
2.7 Å
b
TM1
TM2
TM3
TM4
TM5
TM6
β-sheets NBD1
β-sheets NBD2
TM7
TM8
TM9
TM10
TM11
TM12
Lasso Domain
DHEA-S

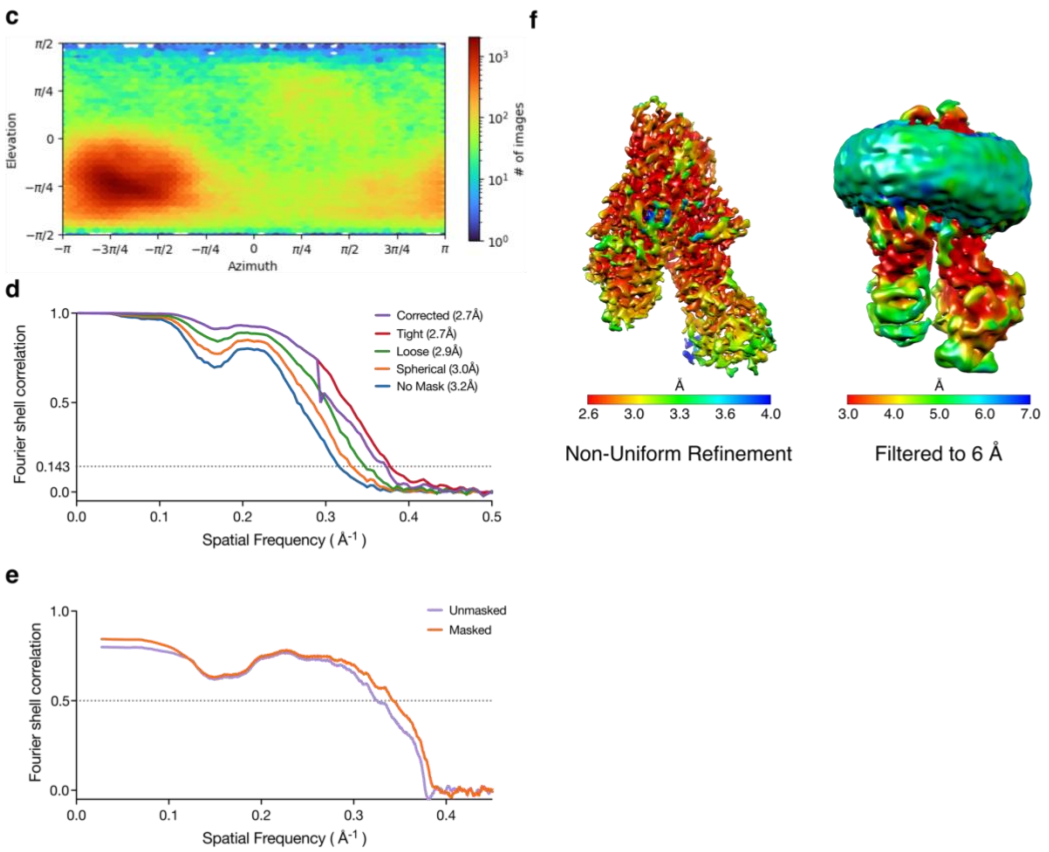


**Supplementary Figure 4. | Image processing of DHEAS-bound MRP4 leading to a 2.7 Å cryo-EM reconstruction.** Cryo-EM analysis of DHEAS-bound MRP4 in lipid nanodiscs. **a**, Processing pipeline for DHEAS-bound MRP4. **b**, Cryo-EM density of the sharpened map overlaid with the model from selected regions. **c**, Euler angle heat-maps for the final DHEAS-bound MRP4 refinement **d**, Gold-standard Fourier shell correlation threshold from the final DHEAS-bound MRP4 refinement. Average resolution determined by using a correlation threshold of 0.143. **e**, Map to model FSC plot indicating correspondence of DHEAS-bound MRP4 atomic model to final density map. **f**, Final density from cryoSPARC refinement colored by local resolution. To show lower resolution features, including NBD1, the map is also shown low-pass filtered to 6 Å.

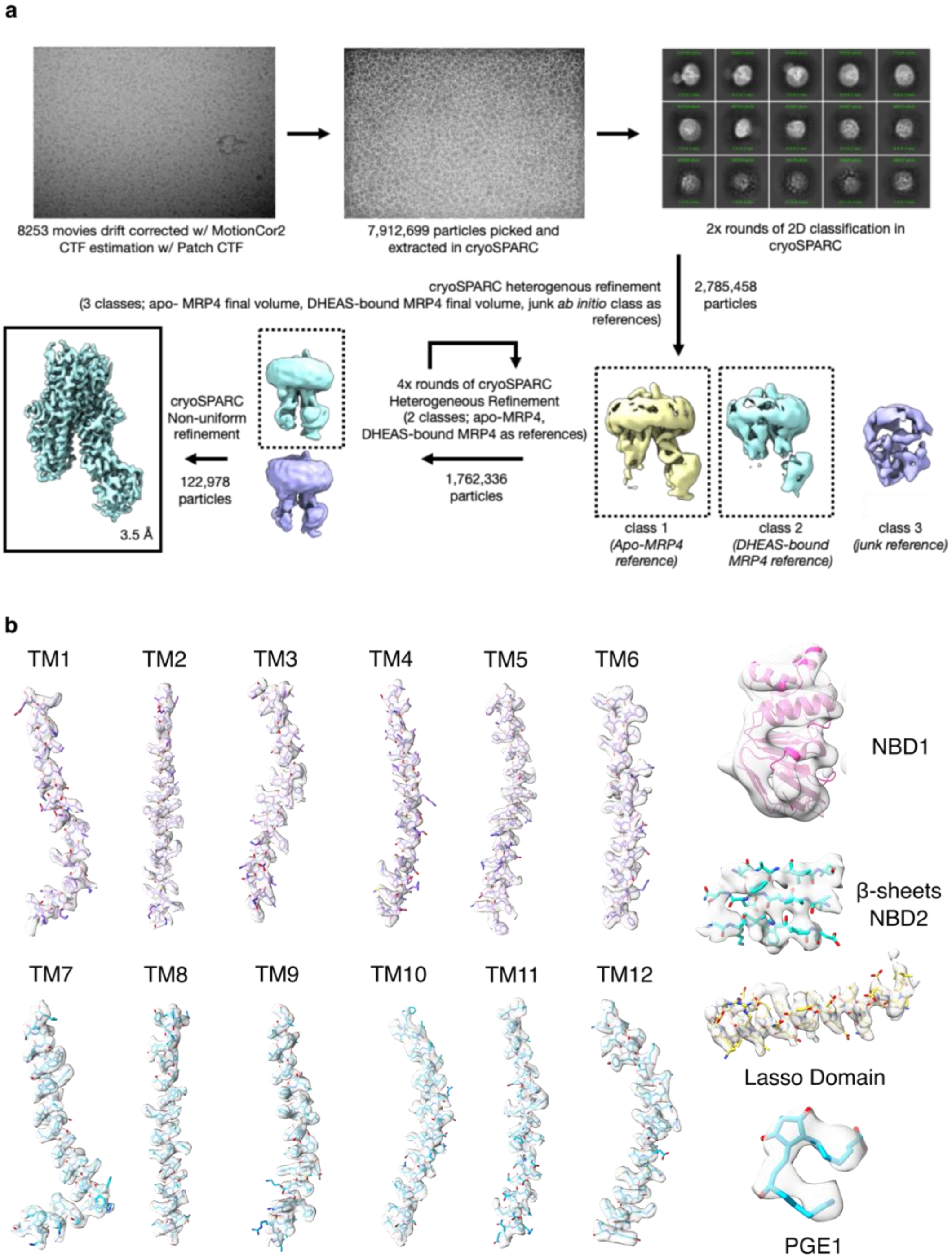
a
8253 movies drift corrected w/ MotionCor2
CTF estimation w/ Patch CTF
7,912,699 particles picked and
extracted in cryoSPARC
2x rounds of 2D classification in
cryoSPARC
cryoSPARC heterogenous refinement
(3 classes; apo- MRP4 final volume, DHEAS-bound MRP4 final volume, junk ab initio class as
references)
2,785,458
particles
4x rounds of cryoSPARC
Heterogeneous Refinement
(2 classes; apo-MRP4,
DHEAS-bound MRP4 as references)
1,762,336
particles
cryoSPARC
Non-uniform
refinement
122,978
particles
3.5 Å
class 1
(Apo-MRP4
reference)
class 2
(DHEAS-bound
MRP4 reference)
class 3
(junk reference)
b
TM1
TM2
TM3
TM4
TM5
TM6
NBD1
β-sheets
NBD2
TM7
TM8
TM9
TM10
TM11
TM12
Lasso Domain
PGE1

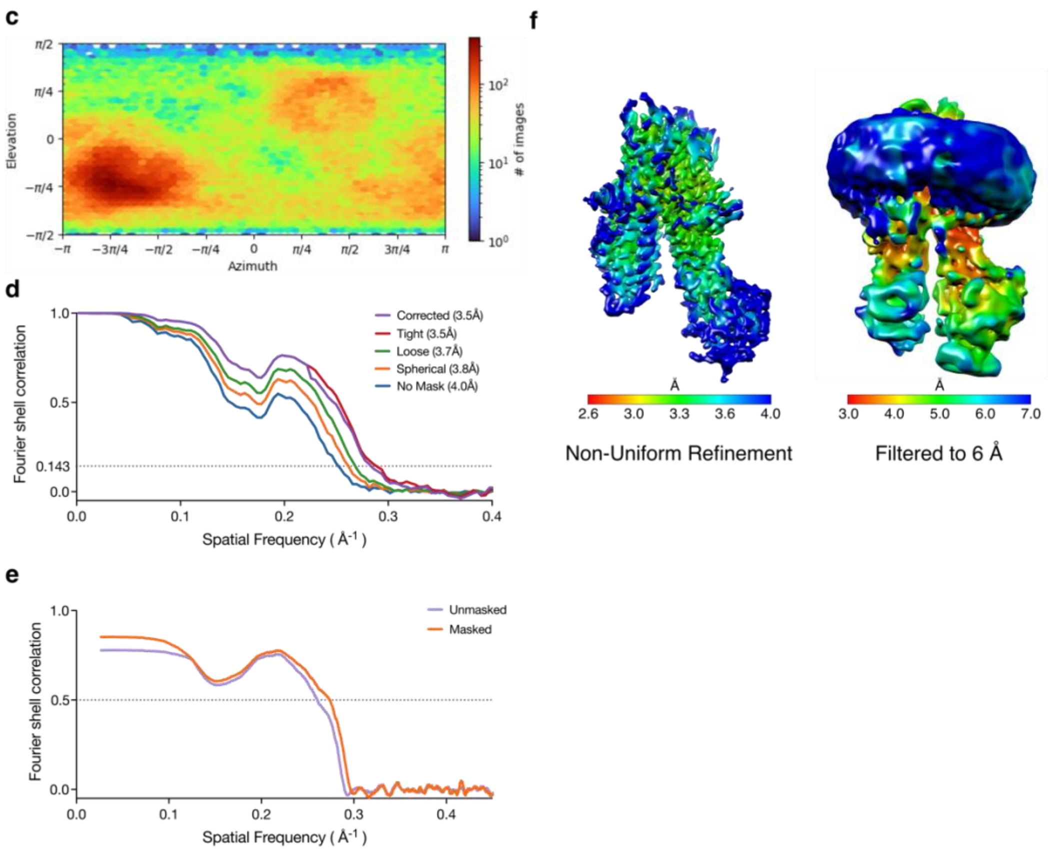


**Supplementary Figure 5. | Image processing of PGE1-bound MRP4 leading to a 3.5 Å cryo-EM reconstruction.** Cryo-EM analysis of PGE1-bound MRP4 in lipid nanodiscs. **a**, Processing pipeline for PGE1-bound MRP4. **b**, Cryo-EM density of the sharpened map overlaid with the model from selected regions. **c**, Euler angle heat-maps for the final PGE1-bound MRP4 refinement **d**, Gold-standard Fourier shell correlation threshold from the final PGE1-bound MRP4 refinement. Average resolution determined by using a correlation threshold of 0.143. **e**, Map to model FSC plot indicating correspondence of PGE1-bound MRP4 atomic model to final density map. **f**, Final density from cryoSPARC refinement colored by local resolution. To show lower resolution features, including NBD1, the map is also shown low-pass filtered to 6 Å.

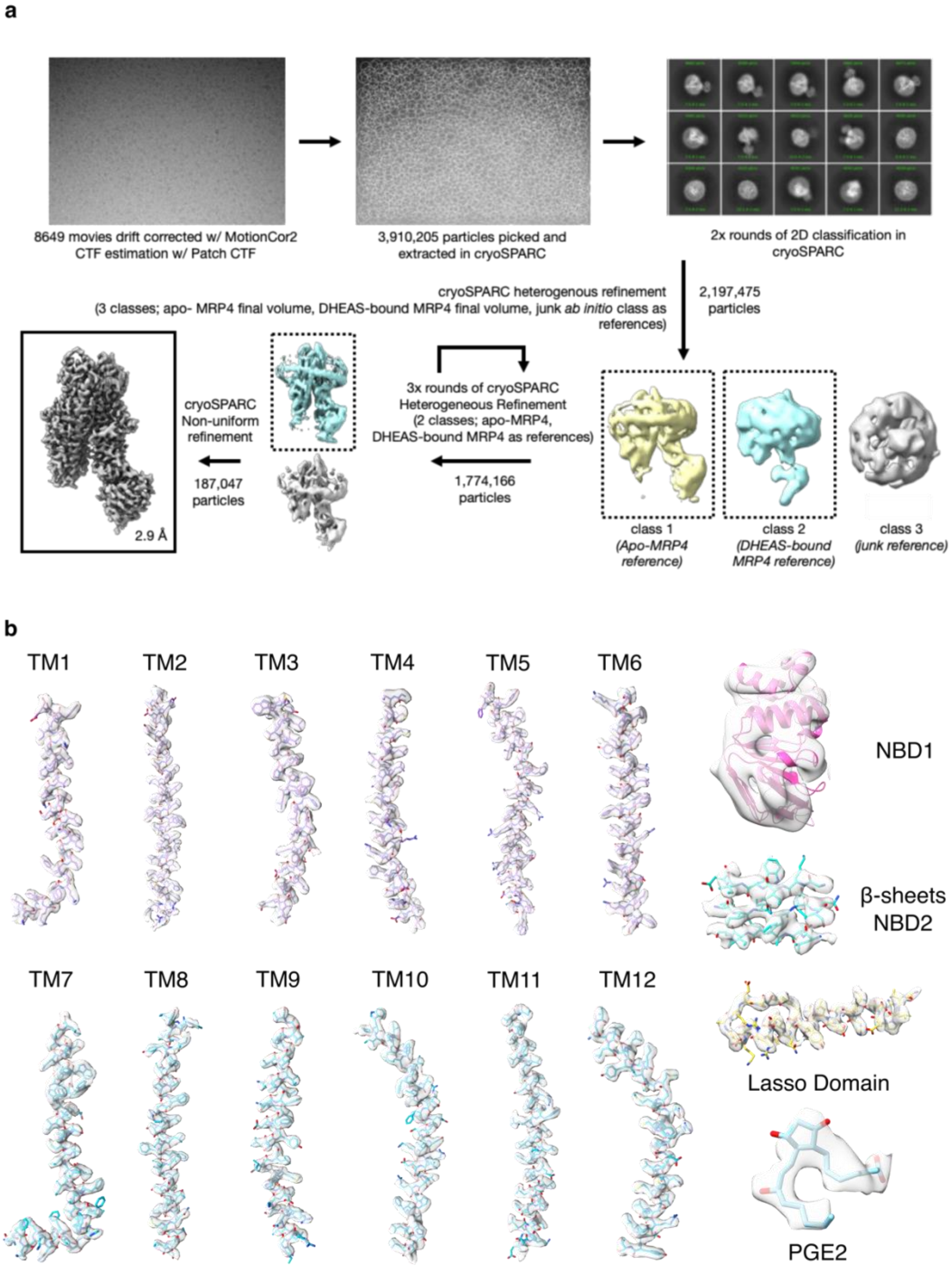
a
8649 movies drift corrected w/ MotionCor2
CTF estimation w/ Patch CTF
3,910,205 particles picked and
extracted in cryoSPARC
2x rounds of 2D classification in
cryoSPARC
cryoSPARC heterogenous refinement
(3 classes; apo- MRP4 final volume, DHEAS-bound MRP4 final volume, junk ab initio class as
references)
2,197,475
particles
3x rounds of cryoSPARC
Heterogeneous Refinement
(2 classes; apo-MRP4,
DHEAS-bound MRP4 as references)
1,774,166
particles
cryoSPARC
Non-uniform
refinement
187,047
particles
2.9 Å
class 1
(Apo-MRP4
reference)
class 2
(DHEAS-bound
MRP4 reference)
class 3
(junk reference)
b
TM1
TM2
TM3
TM4
TM5
TM6
NBD1
β-sheets
NBD2
TM7
TM8
TM9
TM10
TM11
TM12
Lasso Domain
PGE2

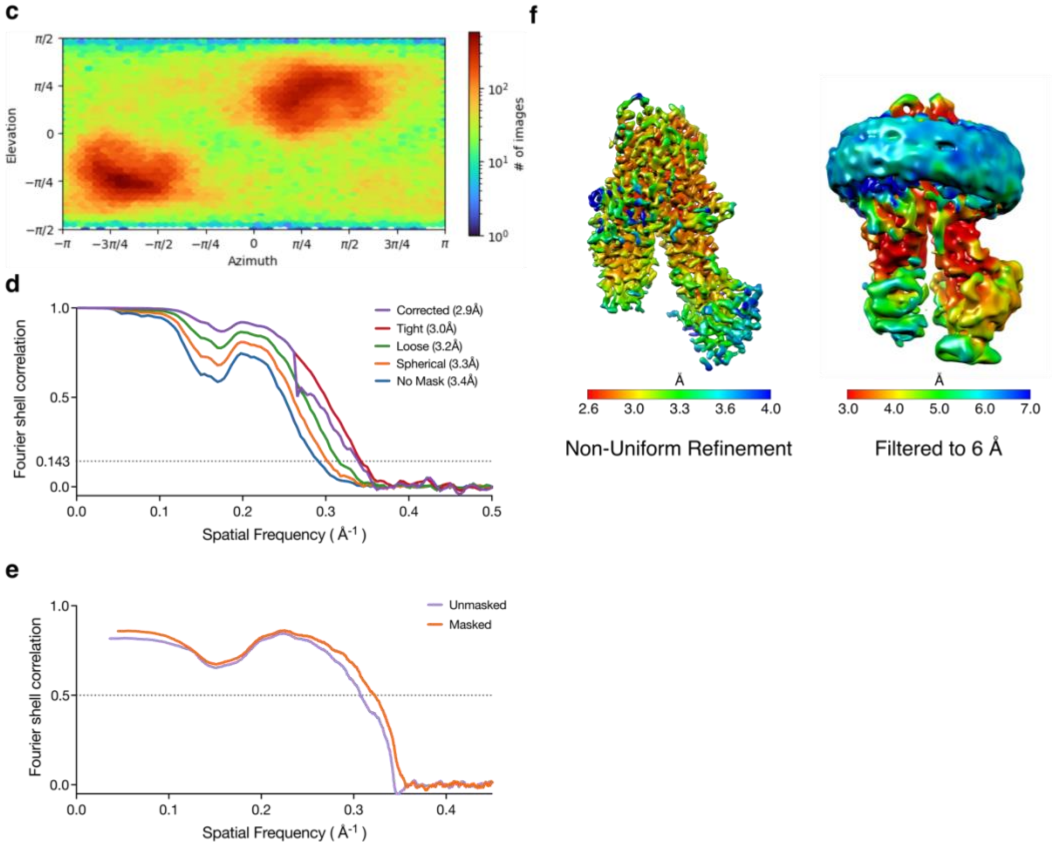


**Supplementary Figure 6. | Image processing of PGE2-bound MRP4 leading to a 2.9 Å cryo-EM reconstruction.** Cryo-EM analysis of PGE2-bound MRP4 in lipid nanodiscs. **a**, Processing pipeline for PGE2-bound MRP4. **b**, Cryo-EM density of the sharpened map overlaid with the model from selected regions. **c**, Euler angle heat-maps for the final PGE2-bound MRP4 refinement **d**, Gold-standard Fourier shell correlation threshold from the final PGE2-bound MRP4 refinement. Average resolution determined by using a correlation threshold of 0.143. **e**, Map to model FSC plot indicating correspondence of PGE1-bound MRP4 atomic model to final density map. **f**, Final density from cryoSPARC refinement colored by local resolution. To show lower resolution features, including NBD1, the map is also shown low-pass filtered to 6 Å.

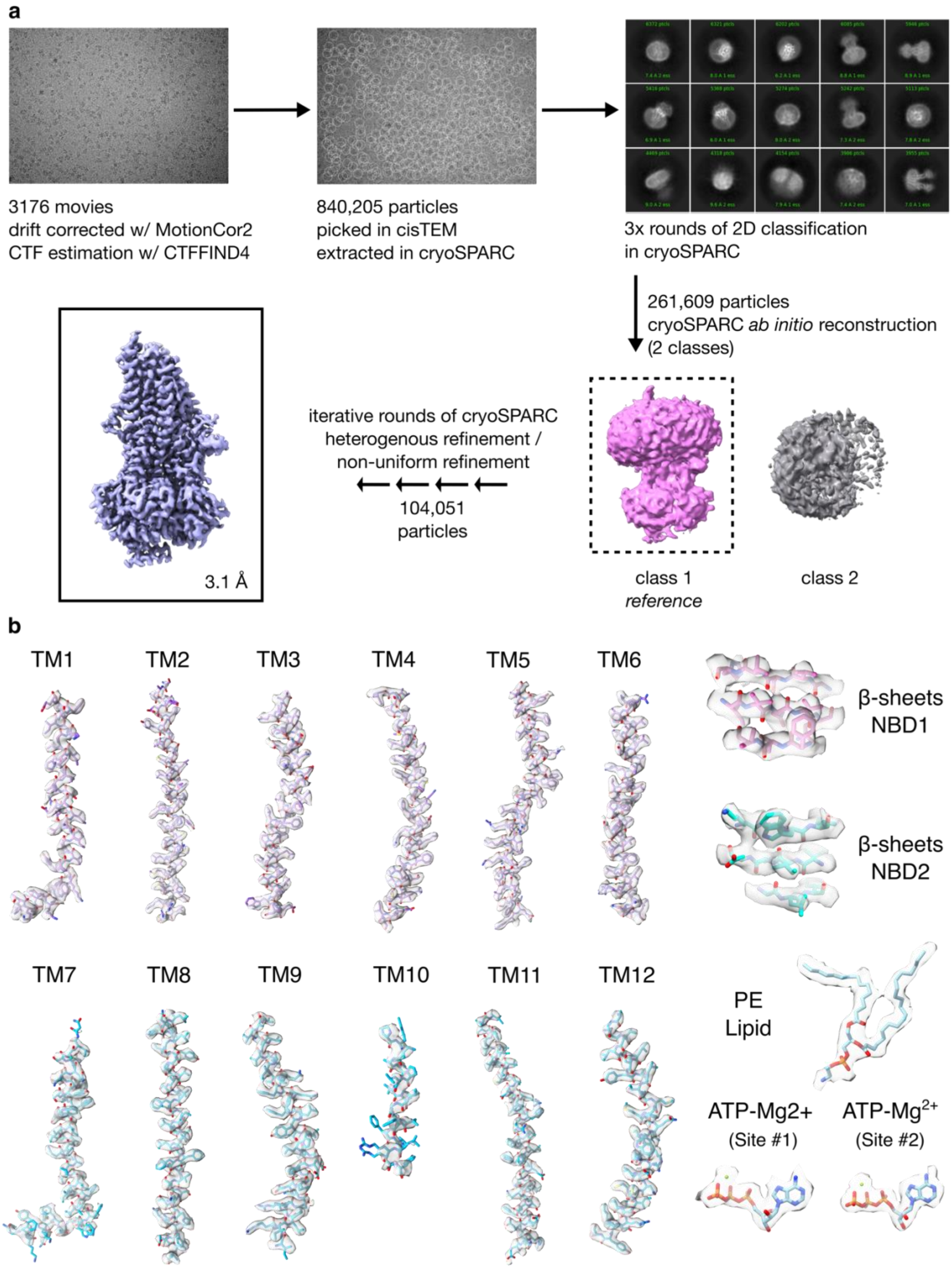
a
3176 movies
drift corrected w/ MotionCor2
CTF estimation w/ CTFFIND4
840,205 particles
picked in cisTEM
extracted in cryoSPARC
3x rounds of 2D classification
in cryoSPARC
261,609 particles
cryoSPARC ab initio reconstruction
(2 classes)
iterative rounds of cryoSPARC
heterogenous refinement /
non-uniform refinement
104,051
particles
3.1 Å
class 1
reference
class 2
b
TM1
TM2
TM3
TM4
TM5
TM6
β-sheets
NBD1
β-sheets
NBD2
TM7
TM8
TM9
TM10
TM11
TM12
PE
Lipid
ATP-Mg2+
(Site #1)
ATP-Mg2+
(Site #2)

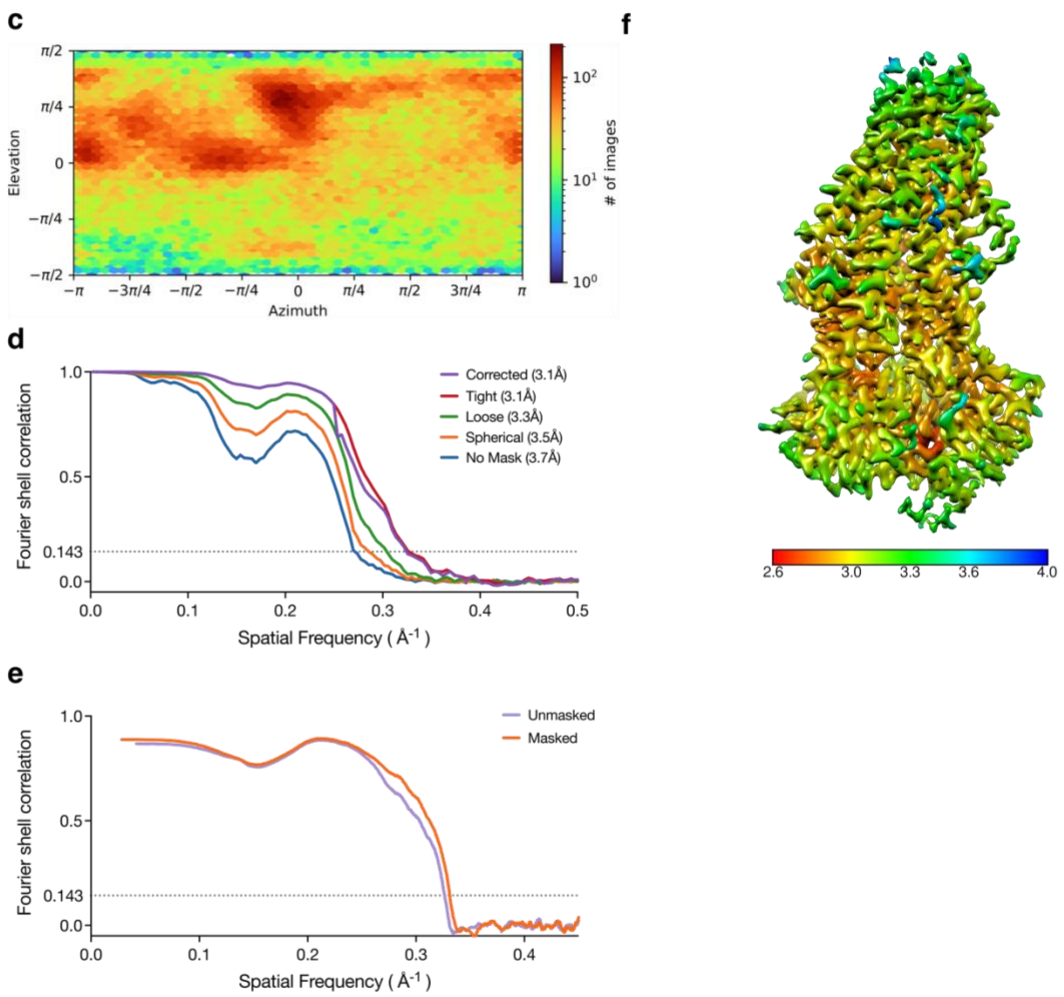


**Supplementary Figure 7. | Image processing of ATP-bound MRP4$_{E1202Q}$ leading to a 3.1 Å cryo-EM reconstruction.** Cryo-EM analysis of ATP-bound MRP4 in lipid nanodiscs. **a**, Processing pipeline for ATP-bound MRP4$_{E1202Q}$. **b**, Cryo-EM density of the sharpened map overlaid with the model from selected regions. **c**, Euler angle heat-maps for the final ATP-bound MRP4 refinement. **d**, Gold-standard Fourier shell correlation threshold from the final ATP-bound MRP4$_{E1202Q}$ refinement. Average resolution determined by using a correlation threshold of 0.143. **e**, Map to model FSC plot indicating correspondence of ATP-bound MRP4$_{E1202Q}$ atomic model to final density map. **f**, Final density from cryoSPARC refinement colored by local resolution.

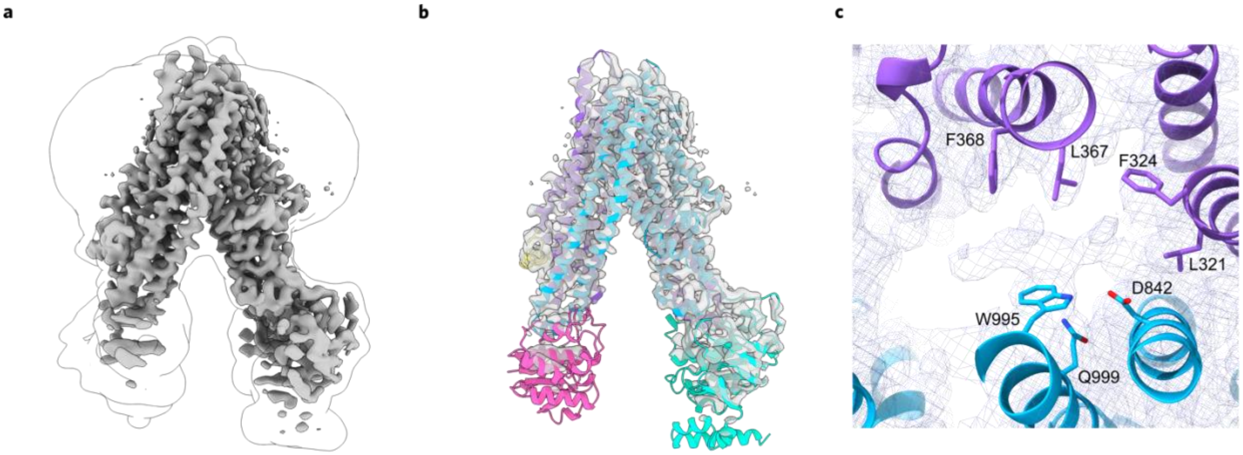


**Supplementary Figure 8. | cAMP does not elicit a conformational change in MRP4 in cryoEM.**

**a** Final density of MRP4 in the presence of 1mM cAMP from cryoSPARC. Sharpened output volume in grey, lowpass filtered volume showing the nanodisc and NBDs as black silhouette.

**b** Rigid body fitting of apo MRP4 into density obtained in the presence of 1mM cAMP. Density in grey, MRP4 domains colored as before. **c** View of substrate binding residues of the rigid-body fit apo MRP4 model in the density obtained for MRP4 in the presence of 1mM cAMP. Map contoured at low threshold, revealing non-protein density similar to our apo MRP4 refinement. Density in purple mesh, MRP4 domains colored as before.

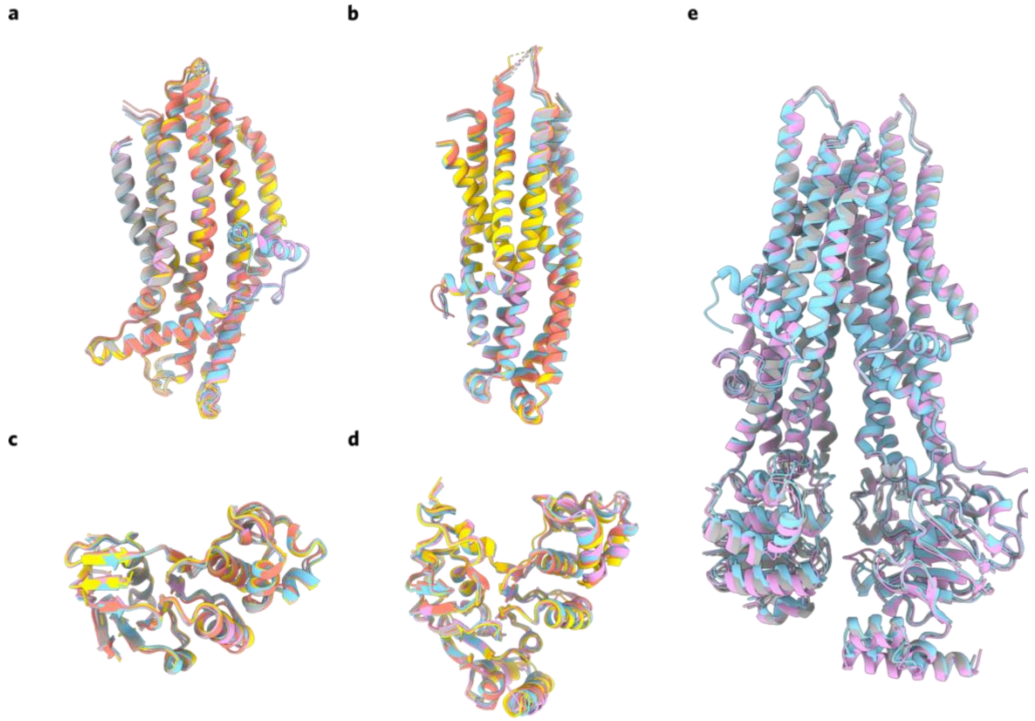


**Supplementary Figure 9. | The domains of MRP4 move as rigid bodies throughout the substrate transport cycle.** Superposition of **a**, bundle 1 (TMs 1, 2, 3, 6, 10, 11) **b**, bundle 2 (TMs 4, 5, 7, 8, 9, 12) **c**, NBD1 and **d**, NBD2 across all five structures. Apo MRP4 in salmon, DHEA-S-bound MRP4 in cyan, PGE-1-bound MRP4 in grey, PGE2-bound MRP4 in pink, ATP-$Mg^{2+}$-bound MRP4 in yellow. **e,** Superposition of the three substrate-bound structures reveals them to share an inward open, narrow conformation. RMSD between all Cα of DHEA-S-bound MRP4 and PGE1-bound MRP4 is 1.2 Å; between DHEA-S-bound and PGE2-bound is 0.87 Å; between PGE1-bound and PGE2-bound is 0.90 Å. Structures are colored as in **a**-**d**.

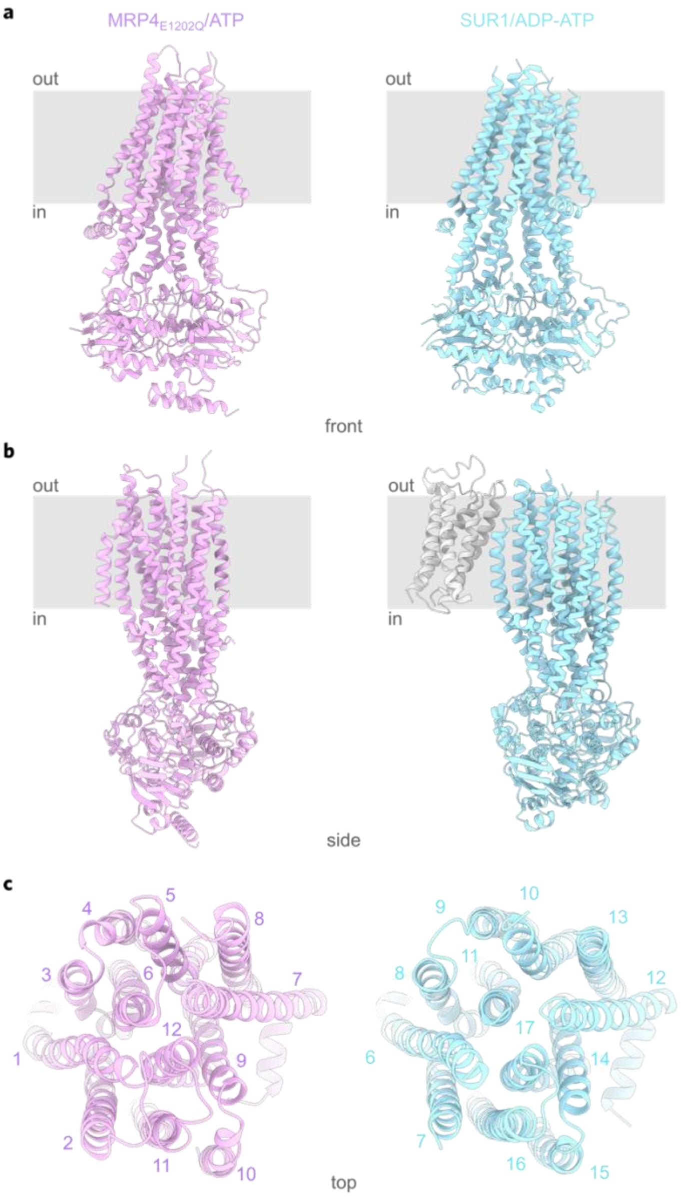


**Supplementary Figure 10. | The outward-facing occluded state of MRP4 closely resembles the structure of SUR1 bound to ADP/ATP. a** Front view, **b** side view, and **c** top view of $MRP4_{E1202Q}$ bound to ATP-$Mg^{2+}$ and SUR1 bound to ADP-ATP (shown as single chain from PDBID:6C3O). $TMD_0$ domain in SUR1 is hidden for clarity in **a** and **c**. Cα RMSD between our ATP-bound structure and SUR1 with $TMD_0$ domain deleted is 4.36 Å. $MRP4_{E1202Q}$ bound to ATP-$Mg^{2+}$ in pink, SUR1 bound to ADP-ATP in cyan, $TMD_0$ domain in light grey.
